# A neurorecording toolkit for longitudinal assessments of transplanted human cortical organoids *in vivo*

**DOI:** 10.64898/2025.12.20.695690

**Authors:** Kate E. Herrema, Elizabeth K. Kharitonova, Daria Bogatova, Madison Wilson, Emily A. Martin, Clara Chung, Francesca Puppo, Shira Klorfeld-Auslender, Pierre Boucher, Li-Huei Tsai, Alysson R. Muotri, Duygu Kuzum, Anna Devor, Timothy M O’Shea, Ella Zeldich, Martin Thunemann

**Affiliations:** Department of Biomedical Engineering, Boston University, Boston, MA, USA; Department of Anatomy and Neurobiology, Boston University School of Medicine, Boston, MA, USA; Department of Biology, Boston University, Boston, MA, USA; Department of Electrical and Computer Engineering, University of California San Diego, La Jolla, CA, USA; Department of Pediatrics, University of California San Diego, School of Medicine, La Jolla, CA, USA; Picower Institute for Learning & Memory, Massachusetts Institute of Technology, Cambridge, MA, USA; Department of Cellular and Molecular Medicine, University of California San Diego, School of Medicine, La Jolla, CA, USA; Center for Academic Research and Training in Anthropogeny, University of California San Diego, La Jolla, CA, USA; Archealization Center, University of California San Diego, La Jolla, CA, USA; Kavli Institute for Brain and Mind, University of California San Diego, La Jolla, CA, USA; Athinoula A. Martinos Center for Biomedical Imaging, Department of Radiology, Harvard Medical School, Massachusetts General Hospital, Charlestown, MA, USA

**Keywords:** brain organoids, neuronal maturation, cranial window, optical microscopy, light sheet

## Abstract

Human cortical organoids (hCOs) are three-dimensional neural cell aggregates that recapitulate certain structural and functional aspects of the developing human cortex. Xenotransplantation of hCOs into the rodent brain enables human-centric modeling of neurodevelopmental processes in a physiologically relevant environment. Here, we present a neurorecording toolkit for longitudinal structural and functional assessment of hCO xenografts as they mature *in vivo*. Single hCOs were implanted into the retrosplenial cortex of adult immunodeficient mice and monitored for up to 8 months. Optical coherence tomography was used for label-free imaging of xenograft vascularization and structure, enabling quantitative assessments of capillary density and graft volume. To probe neuronal activity, human neurons were labeled with a calcium sensor before implantation using either adeno-associated or lentivirus for sparse or dense neuronal labeling, respectively. Fluorescent imaging was conducted using two-photon, widefield, and swept confocally-aligned planar excitation microscopy for single cell, whole-graft, and volumetric calcium imaging, respectively. Results from these modalities indicate an increase in neuronal activity and synchronicity over time during *in vivo* graft maturation. Further, we chronically implanted surface graphene microelectrode arrays (gMEAs) and performed recordings of xenograft and host local field potential signals simultaneously with 2P calcium imaging, confirming the spatial localization and human origin of electrical signals recorded at the xenograft surface.

## Introduction

Over the past decade, human induced pluripotent stem cell (hiPSC)-derived cortical organoids (hCOs) have emerged as powerful, three-dimensional neural cell culture systems that recapitulate certain structural and functional aspects of the developing human cortex to model cortical development and pathologies *in vitro* (Kadoshima *et al*. 2013, Lancaster *et al*. 2013, Pasca *et al*. 2015, Birey *et al*. 2017, Madhavan *et al*. 2018, Qian *et al*. 2019, Trujillo *et al*. 2019, Li *et al*. 2022). Compared to two-dimensional cell culture models, hCOs display increased cellular diversity, more physiologically relevant cytoarchitectures, and more complex intercellular interactions. Furthermore, hCOs are a valuable alternative to animal models as they recapitulate aspects of the transcriptomic, epigenetic, and electrophysiological profile of human fetal brain development in mid-gestation (Camp *et al*. 2015, Amiri *et al*. 2018, Trujillo *et al*. 2019, Gordon *et al*. 2021) and preserve some human-specific genetic traits and phenotypes associated with neurodevelopmental and neurodegenerative aberrations (Parr *et al*. 2017). Despite these advantages, hCOs have several key limitations. First, without vasculature, hCOs suffer from a lack of oxygen and nutrient perfusion which limits their growth and leads to the formation of a necrotic core during extended *in vitro* culture (Qian *et al*. 2019). Second, hCOs lack important cell populations such as microglia and endothelial cells. Lastly, without inputs from functionally relevant neuronal circuits or the presence of natural synaptic targets, hCOs are unable to form more mature neuronal networks. To address these challenges, attention has turned to xenotransplantation models.

Xenotransplantation of hCOs into the rodent brain facilitates graft vascularization, improved cellular diversity and maturation, and functional integration of hCO-derived neurons into host circuits, enabling human-centric modeling of neurodevelopment and disease with improved physiological relevance (Daviaud *et al*. 2018, Mansour *et al*. 2018, Kitahara *et al*. 2020, Shi *et al*. 2020, Revah *et al*. 2022, Wilson *et al*. 2022, Jgamadze *et al*. 2023, Schafer *et al*. 2023, Kelley *et al*. 2024, Wang *et al*. 2025). Recent studies have demonstrated, primarily using electrophysiological techniques, that xenotransplanted hCO neurons achieve greater functional maturity compared to their *in vitro* counterparts (Revah *et al*. 2022, Wilson *et al*. 2022), but the temporal trajectory of this phenomenon has not been thoroughly characterized. To better track and understand these dynamic processes, there is a need for sophisticated tools to longitudinally monitor the growth and activity of hCO xenografts *in vivo*. Toward this end, we previously established a multimodal monitoring strategy using two-photon (2P) microscopy to assess xenograft vascularization in combination with optically transparent graphene microelectrode arrays (gMEAs) to evaluate electrophysiological activity of the graft and surrounding host cortex (Wilson *et al*. 2022). Using this platform, we longitudinally assessed xenotransplanted hCOs *in vivo* and demonstrated the functional integration of human neurons with mouse neuronal networks (Wilson *et al*. 2022). In the present study, we significantly expanded this platform by incorporating different labeling strategies and combining additional imaging modalities to enable complementary functional, multiscale neuronal imaging and label-free assessments of graft structure and integration.

Functional assessment of human neurons within xenotransplanted hCOs via calcium imaging has been previously demonstrated using 2P microscopy (Mansour *et al*. 2018, Revah *et al*. 2022) which enables analysis of neuronal network dynamics at the single-cell level with high spatiotemporal resolution. Other commonly used modalities for calcium imaging include one-photon (1P) widefield microscopy (Cardin *et al*. 2020) and light-sheet methods compatible with *in vivo* imaging, namely swept confocally-aligned planar excitation (SCAPE) microscopy (Bouchard *et al*. 2015, Voleti *et al*. 2019). These modalities complement 2P calcium imaging by capturing neuronal activity in larger fields of view (1P), though with decreased spatial resolution and limited depth penetration, or at cellular resolution through high-speed, three-dimensional volumetric imaging (SCAPE). However, neither modality has thus far been applied to measure neuronal activity in hCO xenografts. Furthermore, previous studies, including ours, have used 2P imaging to assess xenograft vascularization. This approach requires fluorescent labelling of the blood plasma via injected fluorescent tracers and offers limited contrast between graft and host tissue (Mansour *et al*. 2018, Wilson *et al*. 2022). High-contrast imaging of the hCO host-graft interface has recently been demonstrated using T2-weighted magnetic resonance imaging (MRI) (Revah *et al*. 2022). Here, we demonstrate optical coherence tomography (OCT) as an alternative, label-free imaging modality. OCT has been used in prior studies for to assess brain vasculature (Wang *et al*. 2017, Tang *et al*. 2021, Tang *et al*. 2025) and to longitudinally assess cell graft remodeling in stroke lesions through a cranial window (Adewumi *et al*. 2025).

In this work, we expand our neurorecording toolkit for longitudinal functional and structural assessments of xenotransplanted hCOs *in vivo* by leveraging four complementary imaging modalities: OCT for label-free assessments of xenograft vascularization and structural integration, and 2P, 1P, and SCAPE microscopy for functional calcium imaging of hCO xenograft neurons. To augment functional imaging readouts, we combined 2P calcium imaging with simultaneous electrophysiological measurements of the hCO xenograft using optically transparent surface gMEAs. Together, these tools enable robust, multiscale evaluations of the dynamic structural and functional aspects of hCO integration *in vivo*. Our findings demonstrate that xenotransplanted hCOs mature progressively over time in accordance with known neurodevelopmental milestones, advancing their translational potential for studying human brain development and pathologies.

## Results

### A multimodal neurophotonics platform for longitudinal structural and functional assessments of transplanted hCOs

To study human neural development *in vivo*, we grafted single hCOs into the retrosplenial cortex of adult (12-20 weeks) immunodeficient (NOD/SCID) mice along with an optical window to provide access for optical imaging (Mansour *et al*. 2018, Wilson *et al*. 2022) (**Figure 1A**). A subset of animals received surface gMEAs, which are compatible with optical imaging (Liu *et al*. 2017, Thunemann *et al*. 2018), to monitor electrical activity. An overview of individual subjects and experimental conditions is provided in **Supplementary Table 1**. In this study, cortical organoids were generated from hiPSCs following two established protocols (Pasca *et al*. 2015, Trujillo *et al*. 2019) and cultured *in vitro* for 40-60 days before implantation. While still in culture, hCOs were virally transduced after 25 days *in vitro* (DIV) to induce neuronal expression of a green fluorescent genetically encoded calcium indicator, either GCaMP6s or 8s under the human synapsin promoter (**Figure 1B**), together with a red structural fluorophore (tdTomato or mScarlet under a general promoter). Following implantation, grafted animals were monitored for up to 8 months (**Supplementary Figure 1**), after which they were sacrificed for post-mortem analyses. After implantation, xenotransplanted hCOs were visible under brightfield microscopy (**Figure 1C**). Successful engraftment of hCOs derived from either protocol was achieved and verified by post-mortem immunostaining for human cells with the STEM121 antibody (**Figure 1D, Supplementary Figure 2**). To monitor xenotransplanted hCO structure and functional maturation longitudinally, we used OCT for label-free assessments of hCO vascularization, volume, and gross morphology as well as 2P, 1P, and SCAPE microscopy for single-cell, widefield, and volumetric calcium imaging in human neurons, optionally combined with simultaneous electrocorticography (ECoG) recordings using surface gMEAs (**Figure 1E**).

**Figure 1.**
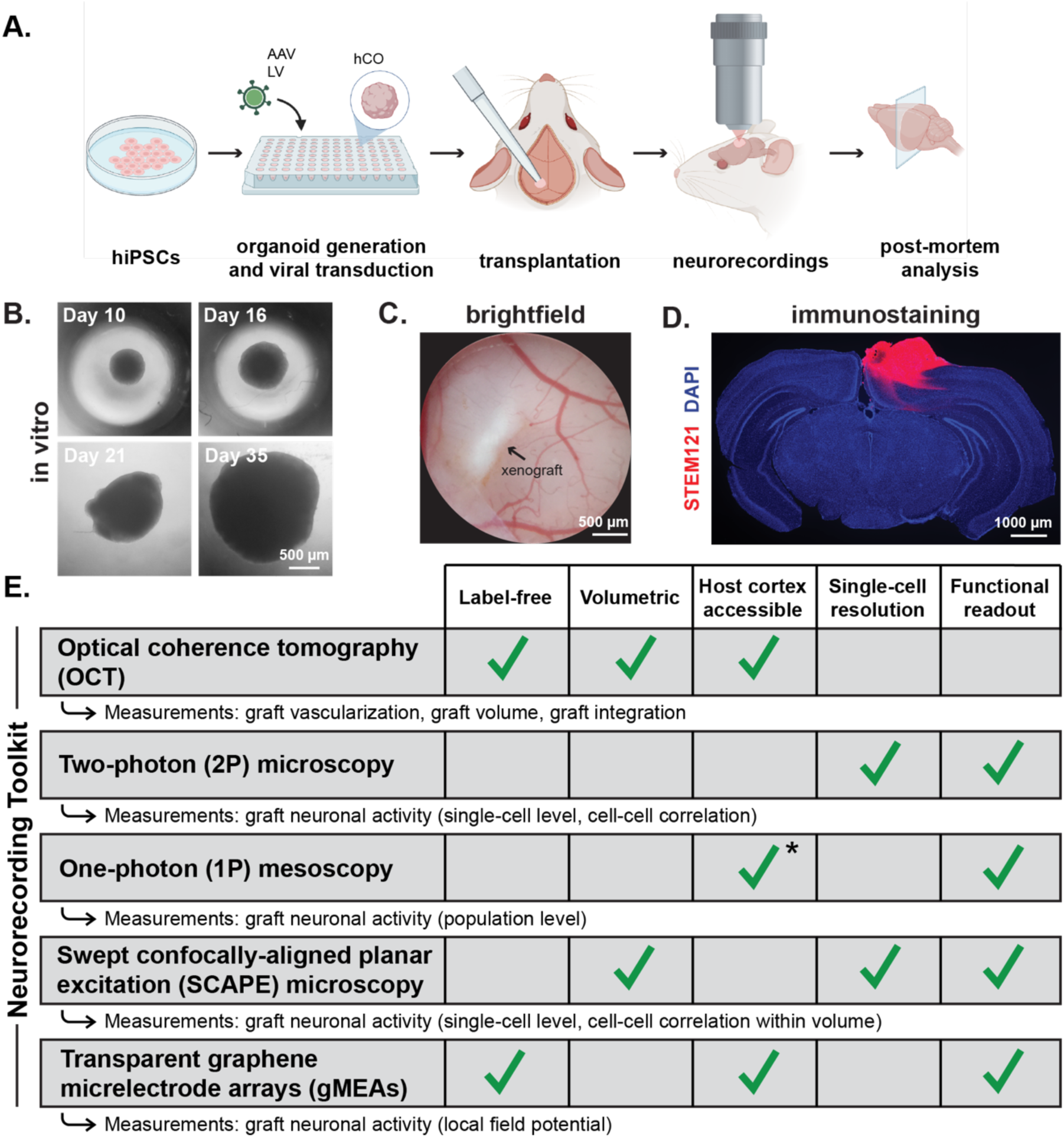
Overview of our *in vivo* neurorecording toolkit for longitudinal assessments of transplanted human cortical organoids (hCOs). A. Schematic of experiment workflow for generation, transduction, transplantation, and assessment of hCOs. Schematic created in <u>BioRender</u>. Abbreviations: *hIPSC*, human-induced pluripotent stem cell, *hCO*, human cortical organoid, *AAV*, adeno-associated virus, *LV*, lentivirus. B. Brightfield image of hCOs (here: Protocol B) during *in-vitro* culture and viral transduction prior to transplantation. C. Brightfield image of transplanted hCO in the retrosplenial cortex of an adult mouse. The image was taken ∼60 days after xenotransplantation. D. Immunofluorescence staining for STEM121 (human cytoplasm) and DAPI (cell nuclei) in a coronal tissue section showing hCO engraftment eight months after xenotransplantation. E. Overview of neurorecording tools used for this study. Asterisk: labeling of host or transgenic host animals is required (not tested in this study).

### OCT allows for label-free, longitudinal structural assessments of graft vascularization and delineation of the host-graft interface

Vascularization of the xenografted hCO is a dynamic process that serves as an important indication of graft health and integration. Previously, we assessed xenograft vascularization using 2P imaging after injecting fluorescent tracers (Wilson *et al*. 2022). Here, we explored alternative, label-free vascular imaging methods. OCT is such a label-free optical imaging technique that has previously been demonstrated for the imaging of brain vasculature, including in cell grafts, by leveraging the optical scattering properties of red blood cells (Wang *et al*. 2017, Tang *et al*. 2021, Adewumi *et al*. 2025). We hypothesized that OCT imaging of hCO xenografts would reveal vascular structures without the use of exogenous contrast agents. Additionally, based on recent work demonstrating the use of OCT for longitudinal assessment of cell grafts in mouse cortical stroke lesions (Adewumi *et al*. 2025), we hypothesized that OCT can be used to delineate the host-graft interface using endogenous tissue contrast, allowing for longitudinal assessment of graft volume and placement.

Mice underwent OCT imaging starting one month after hCO transplantation; angiograms were generated using previously described methods (Wang *et al*. 2017) (**Figure 2A**). In parallel, we performed 2P imaging of blood vessels with Alexa Fluor 680 conjugated to 0.5-MDa dextran (Yao *et al*. 2023) as intravascular tracer (**Figure 2B**). Our results demonstrate a strong agreement between the two modalities (**Figure 2A-C**), justifying the use of OCT as an alternative, label-free method to access graft vascularization. Using OCT, we assessed graft vascularization at monthly intervals, revealing a gradual increase in capillary density over time (**Figure 2D, Supplementary Figure 3)**.

**Figure 2.**
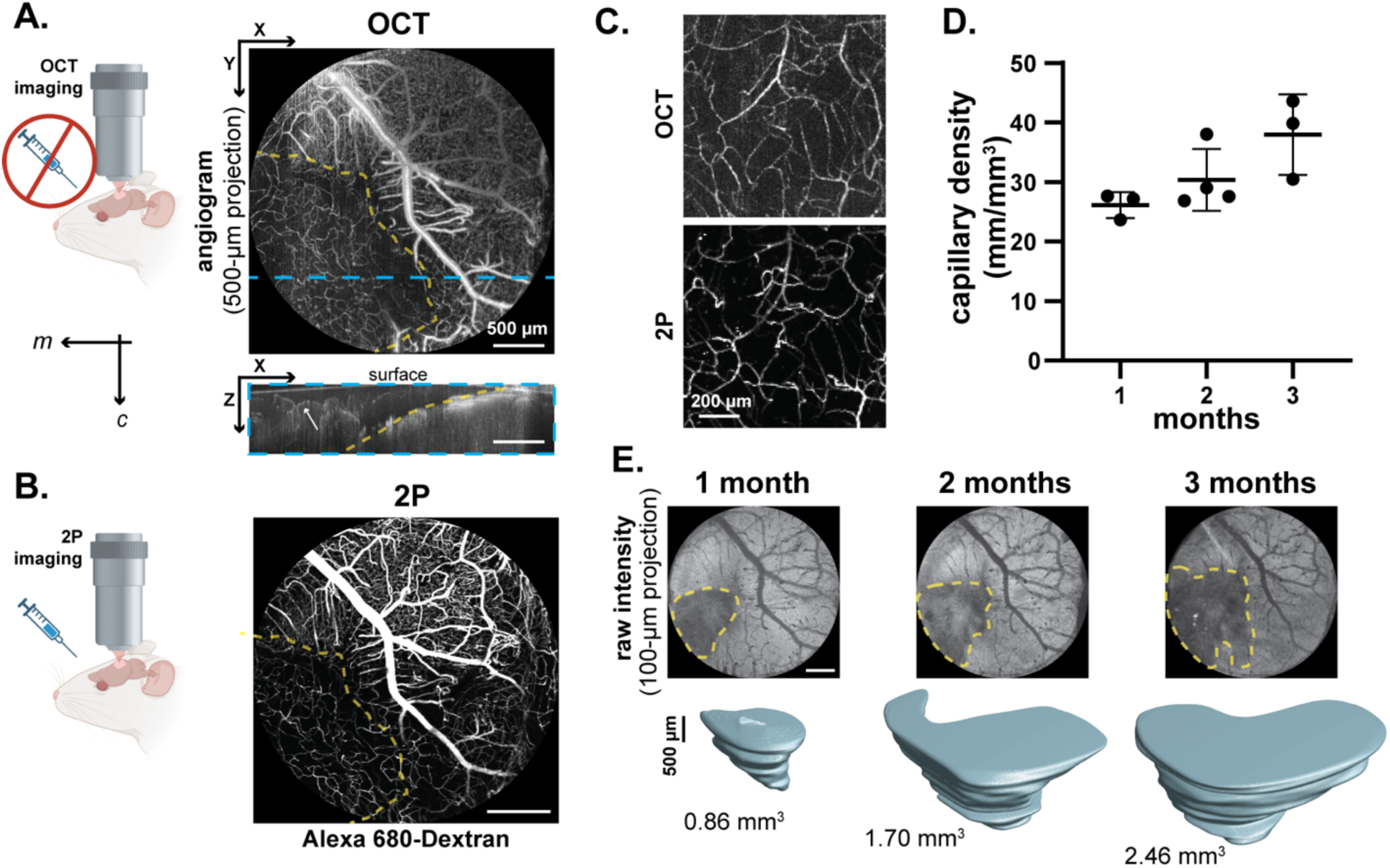
Optical coherence tomography (OCT) enables label-free imaging of xenograft vascularization and morphology. A. Angiogram generated with OCT imaging two months post-transplantation shown as (top) 500-µm maximum intensity projection along the Z axis (MIP_Z_) and (bottom) as 100-µm maximum intensity projection along the X axis (MIP_X_). The yellow dashed line indicates xenograft borders with the host cortex. The blue dashed line indicates the image segment corresponding to the bottom MIP_X_ image. Schematic created in <u>BioRender</u>. Abbreviations: *m*, medial, *c*, caudal. B. Angiogram generated with two-photon (2P) imaging after intravenous injection of fluorescent Alexa-680 Dextran two months post-transplantation for the same animal as shown in panel A. The yellow dashed line indicates xenograft borders with the host cortex. C. Magnified 500-µm MIP_Z_ angiograms showing capillaries within the xenograft as captured by OCT (top) and 2P microscopy (bottom) demonstrating strong agreement between the two modalities. D. Quantification of capillary density within the xenograft using 1 mm x 1 mm x 500 µm MIP_Z_ OCT images at 1, 2, and 3 months after xenotransplantation. The plot shows the mean ± s.e.m. for three animals. E. Volumetric analysis of xenograft size 1, 2, and 3 months after xenotransplantation. (top) Logarithm-normalized, raw intensity, 100-µm MIP_Z_ OCT images. The yellow dashed line indicates xenograft borders with the host cortex. (bottom) Manually segmented, 3D representations of the xenograft and the corresponding estimate of the xenograft volume.

In addition to imaging vascular density, OCT images can be used to delineate graft and host tissue due to endogenous contrast caused by differences in the refractive indices between the two tissues, likely due to variations in myelin content and overall tissue composition (Madhavan *et al*. 2018). We performed image segmentation to extract volumetric information about the graft and were able to assess overall graft morphology and total volume over time. We observed a near three-fold increase in graft volume between one and three months after implantation in the example shown (**Figure 2E**). This indicates that transplanted hCOs undergo large changes in graft volume in the chronic phase following implantation. Notably, while the host-graft interface could be delineated via OCT one month post-implantation in all (N=3) xenografts that were imaged, most xenografts exceeded, at least partially, the bounds of the optical window two months post-implantation, preventing further quantification of the graft volume (**Supplementary Figure 4**).

### Lentiviral transduction yields denser hCO xenograft labelling compared to transduction with adeno-associated viral vectors

To perform longitudinal recordings of neuronal activity via calcium imaging and to assess neuronal maturation in the graft, hCOs were virally transduced in culture before implantation with constructs for expression of GCaMP6s or 8s under control of the human synapsin 1 (hSyn1) promoter. Previously, we and others performed calcium imaging in cultured hCOs after transduction with adeno-associated virus (AAV) (Watanabe *et al*. 2017, Samarasinghe *et al*. 2021). This method successfully labeled human neurons in hCOs *in vitro*, and we were able to visualize neuronal activity at sufficient sampling density at 120 DIV (**Supplementary Video 1**). In contrast, AAV-labeled hCO xenografts showed sparse labeling after transplantation (**Figure 3A**). In line with other studies, we then used lentiviral (LV) vectors to transduce hCOs in culture before transplantation (Revah *et al*. 2022, Kelley *et al*. 2024). To directly compare AAV- and LV-driven GCaMP8s expression in the grafts, hCOs were transduced at 25 DIV with either vector. Following hCO xenotransplantation, GCaMP8s expression and functional activity were assessed using 2P imaging. One month after xenotransplantation, there was a striking difference in the labeling efficiency of AAV-labeled xenografts compared to their LV-labeled counterparts. Additionally, AAV-labeled hCOs exhibited a reduction in labeled neurons *in vivo* from one to three months post-transplantation while LV-labeled hCOs maintained consistent labeling over time (**Figure 3A**). Post-mortem immunostaining 3 months after xenotransplantation revealed that the AAV-hSyn-GCaMP8s construct labeled 0.1% of human cells *in vivo* while the LV-hSyn-GCaMP8s construct was significantly more effective, labeling 11.5% of human cells (p=0.0002; **Figure 3B-C**). Despite differences in labelling efficiency between the two viral constructs, both conditions permitted functional calcium recordings of individual neurons (**Figure 3D**). To validate that labeling differences did not arise from any toxic effects of viral transduction, we performed neuronal (NeuN) staining which showed no significant difference in the density of NeuN-positive cells between AAV- and LV-labeled xenografts (**Supplementary Figure 5**). Thus, we attributed the difference in labeling to mechanistic differences between the two viral vectors (Zheng *et al*. 2018, Fischer *et al*. 2019).

**Figure 3.**
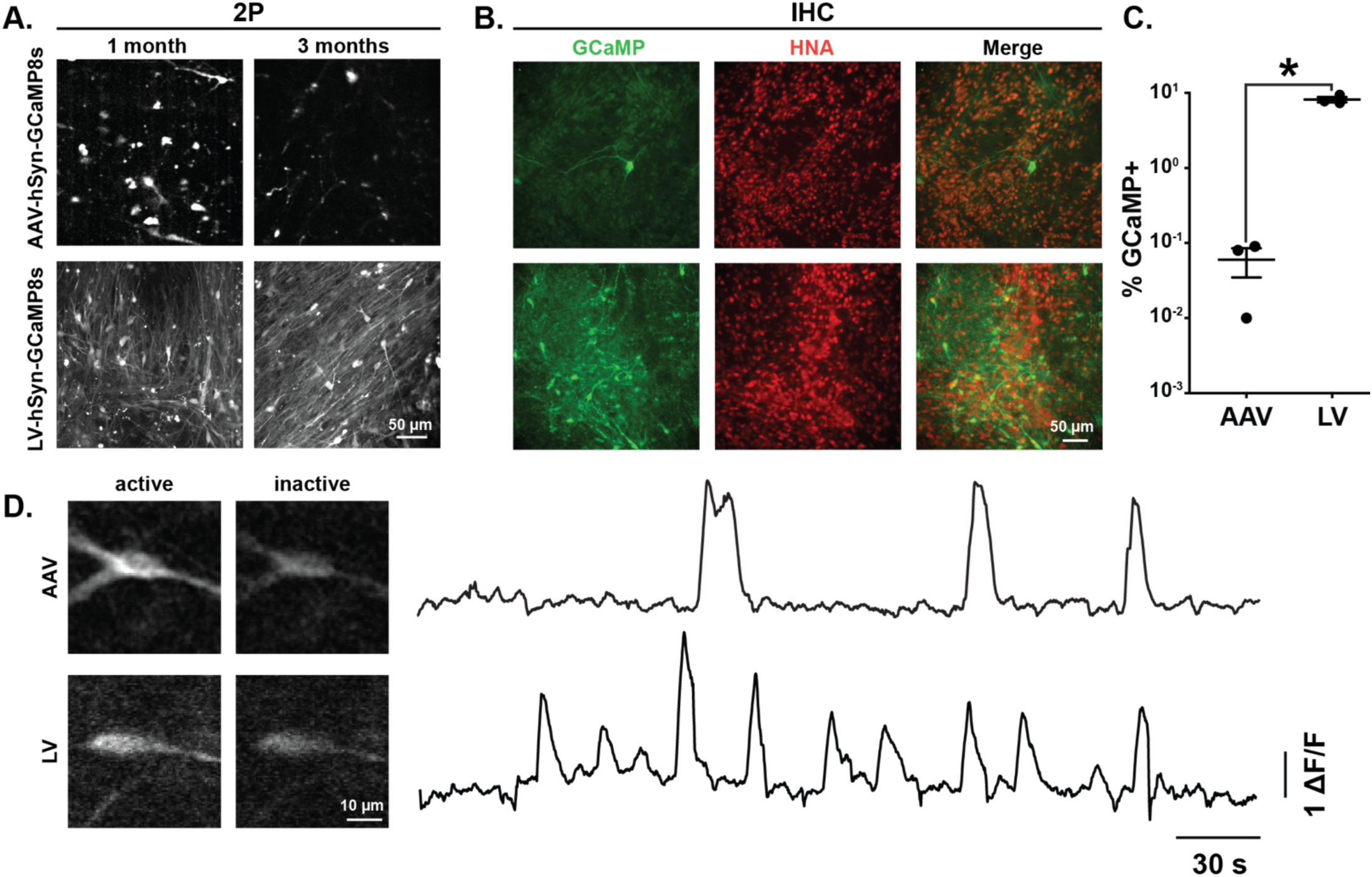
Transduction of human cortical organoids (hCOs) with adeno-associated or lentiviral vectors (AAV or LV, respectively) lead to different densities of GCaMP-expressing neurons after xenotransplantation. A. Two-photon (2P) images from xenografts labelled by either AAV-hSyn-GCaMP8s (top) or LV-hSyn-GCaMP8s (bottom) in live mice one and three months after xenotransplantation. Images were generated from standard deviations of 3-minute time-series recordings. B. Immunohistochemical (IHC) staining for GCaMP-expressing cells with a GFP antibody and human cells with a human nuclear antigen (HNA) antibody. The tissue was isolated three months after xenotransplantation. C. IHC-based quantification of GCaMP labelling density between LV and AAV-labelled xenografts. The bar chart shows mean ± s.e.m. for N = 3 mice per condition. *, P = 0.0002 (two-tailed t-test). D. Representative calcium traces (shown as baseline normalized ΔF/F) recorded with 2P imaging of GCaMP8s-expressing human neurons within the xenograft; hCOs were transduced with AAV (top) or LV (bottom) before transplantation. Animals were recorded while being awake and head-fixed without presentation of external stimuli.

### Longitudinal 2P calcium imaging reveals changes in activity patterns in xenografted human neuron populations

Using AAV or LV to achieve sparse or dense labelling of hCO neurons, respectively, calcium imaging of human neurons expressing GCaMP6s or 8s was performed to investigate single-cell and local network activity (**Figure 4A**). Using 2P microscopy, we were able to observe calcium transients in GCaMP-expressing human neurons *in vivo* for up to eight months after implantation (**Figure 4B-C, Supplementary Figure 1, Supplementary Videos 1-2**). To assess longitudinal changes in neuronal activity in a sufficient number of cells, we used the denser labelling paradigm offered by lentiviral transduction with LV-hSyn1-GCaMP8s and LV-EF1-mScarlet and performed 2P calcium imaging at four-week intervals. For each recording, fluorescence changes in neuronal cell bodies were extracted as relative change to its baseline fluorescence as ΔF/F. To robustly define periods of activity in individual cells, we classified ΔF/F increase exceeding ten standard deviations across the entire time course as a ‘calcium event’; using this threshold, each trace was binarized into periods of activity (calcium events) or inactivity.

**Figure 4.**
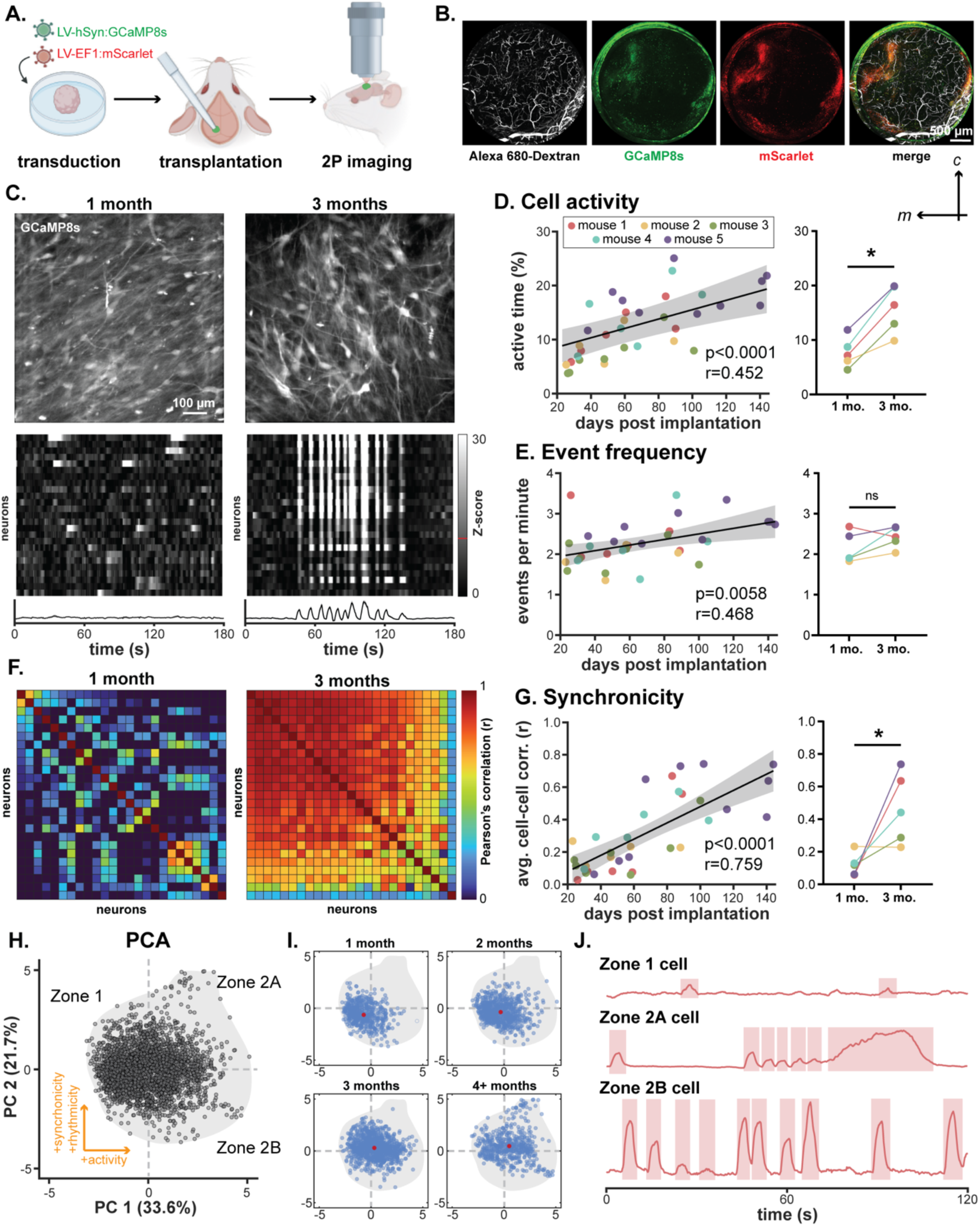
Single-cell activity measured with two-photon (2P) calcium imaging in xenotransplanted human cortical organoids (hCOs) evolves over time. A. Experimental paradigm. For data shown in this figure, hCOs were transduced with LV-hSyn-GCaMP8s and LV-EF1-mScarlet before transplantation; animals were recorded while being awake and head-fixed without presentation of external stimuli. Schematic created with <u>BioRender</u>. B. Representative low-magnification 2P images (maximum intensity projections along the Z axis) of the entire optical window showing vasculature (Alexa 680-Dextran, white), GCaMP8s (green), and mScarlet (red). Abbreviations: *m*, medial, *c*, caudal. C. Two-photon images of GCaMP8s-labelled neurons in the xenograft one and three months after xenotransplantation (top) with corresponding Z-scored heat maps of calcium activity during spontaneous activity (middle) and calcium activity traces averaged across all cells (bottom). The red line on the color bar denotes the activity threshold of 10 standard deviations (Z score = 10). D. Longitudinal changes in neuronal activity over three months for each animal and imaging session (left) and averaged for pairwise comparison (right). Activity of individual cells is determined by the percentage of time the calcium trace for a given cell exceeds a threshold of 10 standard deviations (Z score >10). The scatter plot (left) shows linear mixed effects model (LMM; N=5; p<0.001, r=0.452). The scatter plot (right) shows a pairwise comparison (*, p = 0.001) paired t-test. E. Longitudinal changes in event frequency over three months for each animal and imaging session (left) and averaged for pairwise comparison (right). Events for a given cell are defined as a period where the calcium trace exceeds a threshold of 10 standard deviations (Z score >10). Data from 5 animals. The scatter plot (left) shows linear mixed effects model (LMM; N=5; p=0.0058, r=0.468). The scatter plot (right) shows a pairwise comparison (ns = not significant, p=0.088) paired t-test. F. Cell-to-cell correlation of calcium activity calculated as Pearson’s correlation coefficient between individual cells shown in panel C as measurement of synchronicity. G. Longitudinal changes in cell-to-cell synchronicity over three months for each animal and imaging (left) and averaged for pairwise comparison (right). Synchronicity is calculated as average Pearson’s correlation coefficient across all cells for each field of view. Data from 5 animals. The scatter plot (left) shows linear mixed effects model (LMM; p<0.001, r=0.759). The scatter plot (right) shows the pairwise comparison (*, p=0.026) paired t-test. H. Principal component analysis (PCA) computed from individual cells with at least one calcium event during data acquisition. Data was pooled across all imaging trials (5 animals, 144 trials, 3089 cells). Features estimated for each cell include event frequency, rhythmicity (inter-event-interval CV), activity, event length, event height, and synchronicity. PC 1 represents increases in cell activity while PC 2 represents increases in synchronicity and rhythmicity. Zones 1, 2a, and 2b define differential cell activity phenotypes within the dataset. I. PCA plot binned by month after xenotransplantation with centroid shown in red. J. Representative calcium traces for cells within Zones 1, 2a, and 2b.

Over three months *in vivo*, we observed increasing neuronal activity in xenotransplanted hCOs (p<0.0001, r=0.452), and cell activity was significantly higher at three months than at one month (p=0.0013; **Figure 4D**). The frequency of calcium events also increased during this period (p=0.0058, r=0.468), though it did not reach statistical significance in pairwise comparison at 1 and 3 months (p=0.0884; **Figure 4E**). We also observed increases in the fraction of cells with observable calcium events (p=0.0020, r=0.499) as well as the overall length of calcium events (p=0.0042, r=0.468) and a slight decrease in event height over time (p=0.0333, r=0.356; **Supplementary Figure 6**). We also calculated the coefficient of variance (CV) for the inter-event interval (IEI) which had no observable trend over time in our dataset (p=0.455, r=0.129; **Supplementary Figure 6**).

We further observed a substantial increase in network synchronicity from one to three months after hCO transplantation. One month after transplantation, neurons exhibited minimal calcium event synchronization; however, by three months, we observed synchronized calcium events across most neurons in each field of view (**Figure 4F**). The emergence of synchronized calcium events was strongly correlated with time post-hCO implantation (p<0.0001, r=0.759) and was significantly greater at 3 months than at 1 month after implantation (p=0.0259; **Figure 4G**). While most neuronal populations we imaged over the course of this study followed these trends, a subpopulation exhibited divergent patterns of activity. To better define this group of cells, we pooled all active neurons that we previously recorded along with their characteristic features including active time, event frequency, the covariance of inter-event intervals (IEI CV) as ‘rhythmicity’, event length, event height, and synchronization within the associated field of view (n=3089). We then performed principal component analysis (PCA) and grouped neurons into one-month-wide bins (**Figure 4H-I, Supplementary Figure 6**). Using the PCA weights for each feature, we assigned cells to two “Zones”: Zone 1, characterized by low activity, and Zone 2, characterized by high activity. Overall, neurons recorded at earlier stages occupied Zone 1, while the population shifted toward Zone 2 over time, in agreement with the statistics reported above. However, at later stages, neurons in Zone 2 diverged into two subpopulations which we named Zone 2A and Zone 2B. Neurons in Zone 2A represent a larger population characterized by highly synchronous calcium activity with longer, oscillatory events while neurons in Zone 2B are a smaller population characterized by shorter, more sporadic events that are not synchronized to the larger network (**Figure 4J, Supplementary Figure 6**). The presence of divergent activity patterns could indicate that neurons mature to perform distinct roles within the network, especially at more advanced stages of maturation.

### One-photon mesoscopic imaging enables widefield, whole-xenograft assessments of neuronal activity

While 2P microscopy records calcium events at the cellular level and enables robust cell monitoring and cell-to-cell comparisons, conventional 2P microscopy is limited to a relatively small field of view and a single Z plane. To perform calcium imaging within the xenograft across multiple scales and dimensions, we tested additional imaging modalities; for imaging at a mesoscopic, neuronal population-level, we implemented 1P widefield microscopy using our custom-built system which we previously designed for imaging of neuronal activity via fluorescent indicators as well as hemodynamics via oxy- and deoxyhemoglobin absorption (Doran *et al*. 2024) (**Figure 5A-B)**. This allowed us to visualize calcium events across the entire xenograft surface after implantation of hCOs transduced with LV-hSyn1-GCaMP8s and LV-EF1-mScarlet (**Figure 5C-D, Supplemental Video 4**). At this scale, signals are only detectable when a population of neurons exhibits synchronous surges in calcium levels, making 1P microscopy a suitable method for assessing widespread changes in neuronal activity within the xenograft (Cardin *et al*. 2020, Warm *et al*. 2025). In agreement with findings from 2P calcium imaging, results from 1P imaging show no detectable calcium activity in the early stages after hCO implantation (**Supplementary Figure 7**). Subsequently, we began to observe consistent, large-scale calcium events that were distributed sporadically with several events occurring in short succession followed by longer periods of silence (**Figure 5D**). The distribution of IEIs is right-skewed with most IEIs falling into the range of <50 s and several larger IEIs up to 300s (**Figure 5E**). This indicates that spontaneous activity in the xenograft is not uniform. Future studies could use shifts to the IEI distribution as a signature of specific pathologies or perturbations of this model system.

**Figure 5.**
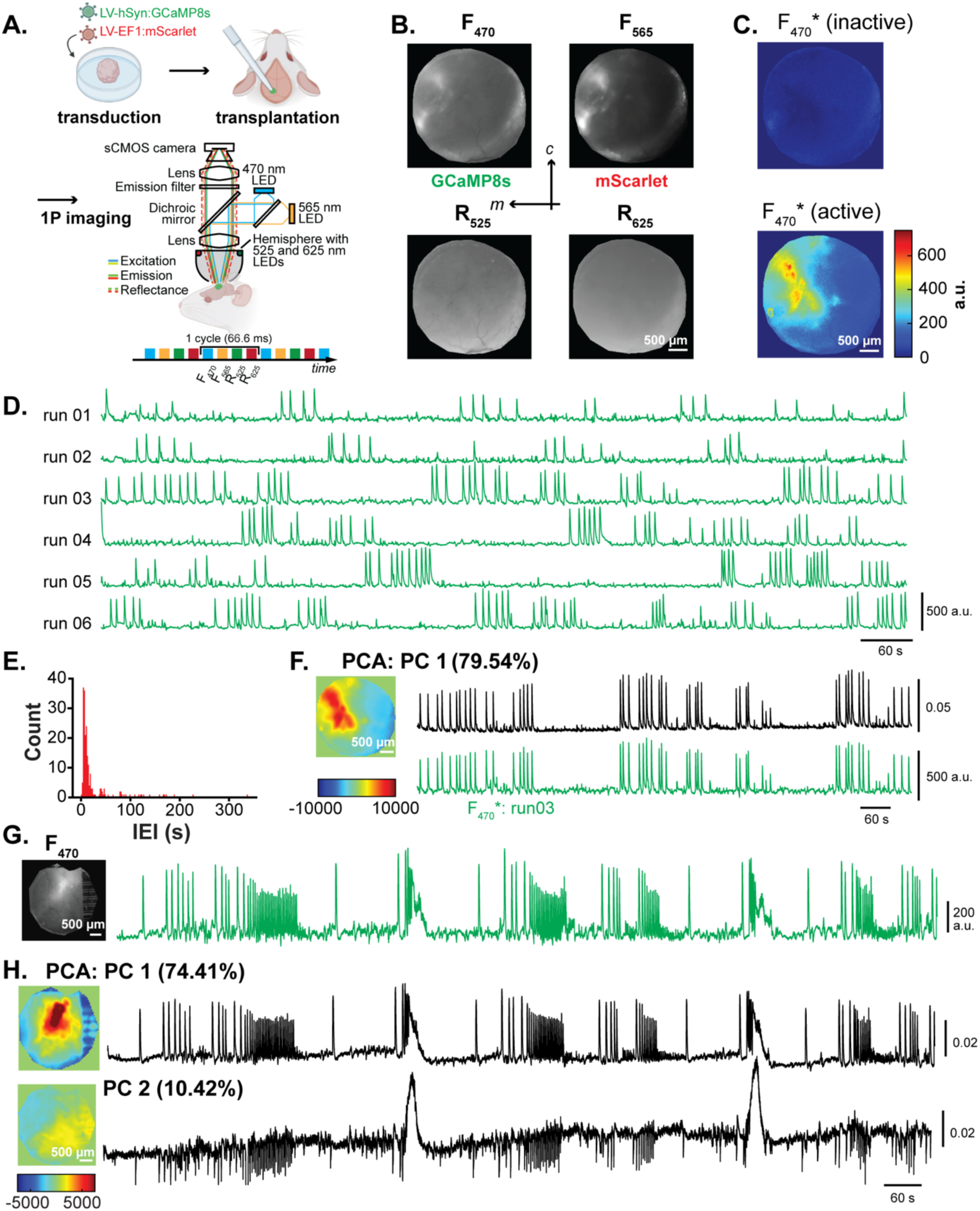
One-photon mesoscale imaging captures whole-graft spontaneous activity. A. Experimental paradigm. For data shown in this figure, hCOs were transduced with LV-hSyn-GCaMP8s and LV-EF1-mScarlet before transplantation. In the widefield microscope, a scientific CMOS camera subsequently records image frames at four different illumination wavelengths at an effective frame rate of 15 Hz; a 565-nm LED (F_565_) excites mScarlet and a 470-nm LED (F_470_) excites GCaMP8s, tissue reflectance is recorded at 525 and 625 nm allowing to estimate changes in hemoglobin concentrations. Microscope schematic adapted from Doran *et al*. (2024). Animals were recorded while awake and head-fixed without presentation of external stimuli. Schematic created partially in <u>BioRender</u>. B. Representative images of an animal with hCO xenograft; the four channels captured by the imaging system are shown: fluorescence at 470-nm and 565-nm excitation (F_470_, GCaMP8s and F_565_, mScarlet), as well as reflectance at 535-nm and 625-nm illumination (R_525_, R_625_). Abbreviations: *m*, medial, *c*, caudal. C. Single frames of the F_470*_ signal during periods of low (top) and high (bottom) calcium activity. To account for artificial signal fluctuations, fluorescence from the F_470_ channel was normalized, pixel-by-pixel, to fluorescence in the F_565_ channel, yielding the normalized F_470_ signal denoted as F_470*_ (see **Methods**). D. Six representative 20-min recordings from the same animal, acquired on different days. Signal intensities in the F470*-channel are derived from manually defined regions of interest outlining the xenograft. E. Histogram of inter-event intervals (IEI) of all calcium events of the recordings shown in panel D. F. Representative results of a pixel-wise principal component analysis (PCA) performed for the F_470_* timeseries of run03 in panel D. Before PCA, data was binned by a factor of 4 in space. A heatmap representing the weight factor across space and the corresponding trace of the first component (PC 1) are shown. For this run, PC 1 accounts for 79.54% of total variability leading to high similarity between the time courses of PC 1 (black) and the corresponding F_470*_ trace (green). G. Temporal average of the F_470*_ timeseries from a second animal and corresponding F_470*_ trace for a 20-min recording. H. Pixel-wise PCA performed on the dataset shown in G; weight maps and time traces for components PC 1 and PC 2 are shown; they account for 74.41% and 10.42% of the total variability, respectively. Note the occurrence of two prolonged events both in PC 1 and PC 2, while shorter events occur mainly in PC 1.

Additionally, we assessed the spatial features of global calcium events in 1P imaging data. While most calcium events occurred globally throughout the graft, we observed some events with a broader or more restricted extent across the xenograft (**Figure 5F**). For further analysis, we reduced the spatial resolution of the data and performed pixel-wise PCA. Results from PCA revealed that even in global, whole-graft calcium events, there are distinct components of the signal that were localized to spatially segregated regions within the xenograft (**Figure 5G-H**). In the example shown, the first principal component (PC 1, 74.41%) is detected throughout the bulk of the xenograft, closely mirroring the original signal. The second component (PC 2, 10.42%) is spatially distinct from PC 1. In contrast to PC 1, PC 2 primarily reflects two longer calcium events that are seen in the original signal while signals in PC 1 are not present in the region defined by PC 2 (**Figure 5H**). These findings suggest that calcium activity is not homogeneous throughout the xenograft but rather reflects region-specific neuronal dynamics.

### SCAPE microscopy enables volumetric calcium imaging in the xenograft

To achieve volumetric imaging of the xenograft to depths of ∼200-250 µm, we used SCAPE, an *in vivo* light sheet microscopy technique based on a single objective for light delivery and detection (Bouchard *et al*. 2015, Voleti *et al*. 2019) (**Figure 6A**). SCAPE enables high-speed, volumetric recordings at single-neuron resolution within the xenograft (**Figure 6B-D**). We hypothesized that this modality would further illuminate the spatial features of calcium signaling in the xenograft by enabling visualization of activity, especially along the depth axis. In a pilot experiment of this modality in one animal, we observed synchronous calcium events in neurons spanning this axis (**Figure 6D**). The synchronous behavior of these cells recorded for four months after hCO xenotransplantation aligns with our observations from 1P and 2P calcium imaging at similar time points. This highlights the capability of SCAPE microscopy to capture the volumetric dynamics of calcium signaling, revealing the spatial relationships of calcium events not accessible to planar imaging modalities.

**Figure 6.**
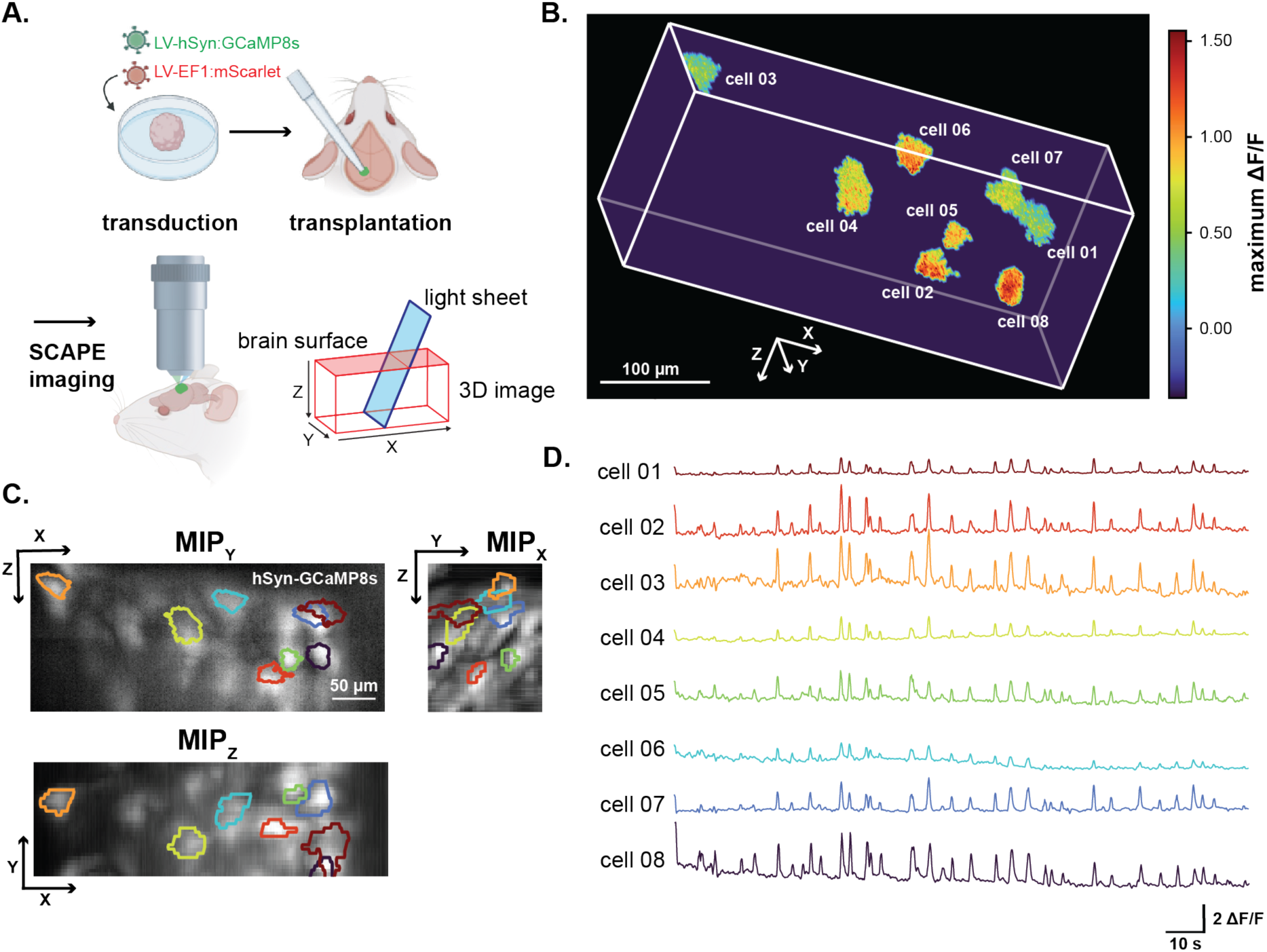
Swept confocally-aligned planar excitation (SCAPE) microscopy permits volumetric imaging of single-neuron calcium events across xenograft depth. A. Schematic of SCAPE imaging experimental paradigm and geometry of the light sheet. Images are captured as the light sheet sweeps across the tissue. Recorded data are reconstructed into a three-dimensional volume recorded at an effective rate of 10 volumes per second. Recording was performed four months after implantation of hCOs labeled with LV-hSyn-GCaMP8s and LV-EF1-mScarlet while the animal was awake and head-fixed without presentation of external stimuli. Panel created partially in <u>BioRender</u>. B. Volume rendering of a single SCAPE volume (131 x 171 x 408 µm) showing segmented regions of interest (ROIs) representing individual cells. Voxels within each extracted ROI are colored based on the maximum ΔF/F (GCaMP8s) within a 136-s recording. C. Maximum intensity projections from the single SCAPE volume shown in panel B along the X, Y, and Z dimensions (MIP_X_, MIP_Y_, and MIP_Z_) of GCaMP8s-expressing human neurons within the xenograft. ROI outlines highlight individual cells. D. Time traces of signal changes (as baseline-normalized ΔF/F) for the ROIs shown in panel B and C. Colors of the individual traces correspond to colors of the cell ROIs in panel C.

### Graphene microelectrode arrays record spatially localized LFP signals corresponding to calcium events in human neurons

In our previous work (Wilson *et al*. 2022), we used optically transparent gMEAs to record local field potentials (LFPs) in xenotransplanted hCOs; however, at that time, our xenografts did not express optical sensors. In this study, we expanded on our previous results by simultaneously recording electrophysiological signals in the xenograft and performing functional calcium imaging using hCOs transduced in culture with AAV-hSyn1-GCaMP6s-P2A-nls-dTomato (**Figure 7A**). Graphene MEAs were implanted over the cortical surface and covered both the xenograft and host cortex (**Figure 7B-C**). Simultaneous recordings of LFP and calcium activity show that discrete calcium transients in individual cells align in space and time with LFP deflections recorded by electrodes near the optical field of view (**Figure 7D-F, Supplementary Figure 8**). Given the electrode surface area and impedance, it is unlikely that only the GCaMP6s-expressing cell(s) within the imaging FOV contribute(s) to the LFP signal detected at the xenograft surface; instead, LFP signals likely represent synchronous surges in the activity of human neurons which aligns with our previous observations that neurons within the xenograft show synchronous activity. Nevertheless, LFP signals recorded with electrodes further away from the imaging FOV show smaller to undetectable LFP deflections while electrodes covering the host cortex also record different patterns of activity (**Figure 7F, Supplementary Figure 8**). Most electrodes exhibited sufficiently low impedance for recording LFPs and remained stable over time for up to eight months in vivo (**Figure 7G**); however, any electrodes with impedances measured greater than 5 MΩ were excluded from further analyses. Sudden increases in the impedance of individual electrodes were likely caused by failure of the connection between array and recording equipment after repeated connection cycles. These results demonstrate that electrical signals originating from the xenograft are spatially confined to electrodes covering it, enabling simultaneous assessment of host and xenograft activity at high temporal resolution.

**Figure 7.**
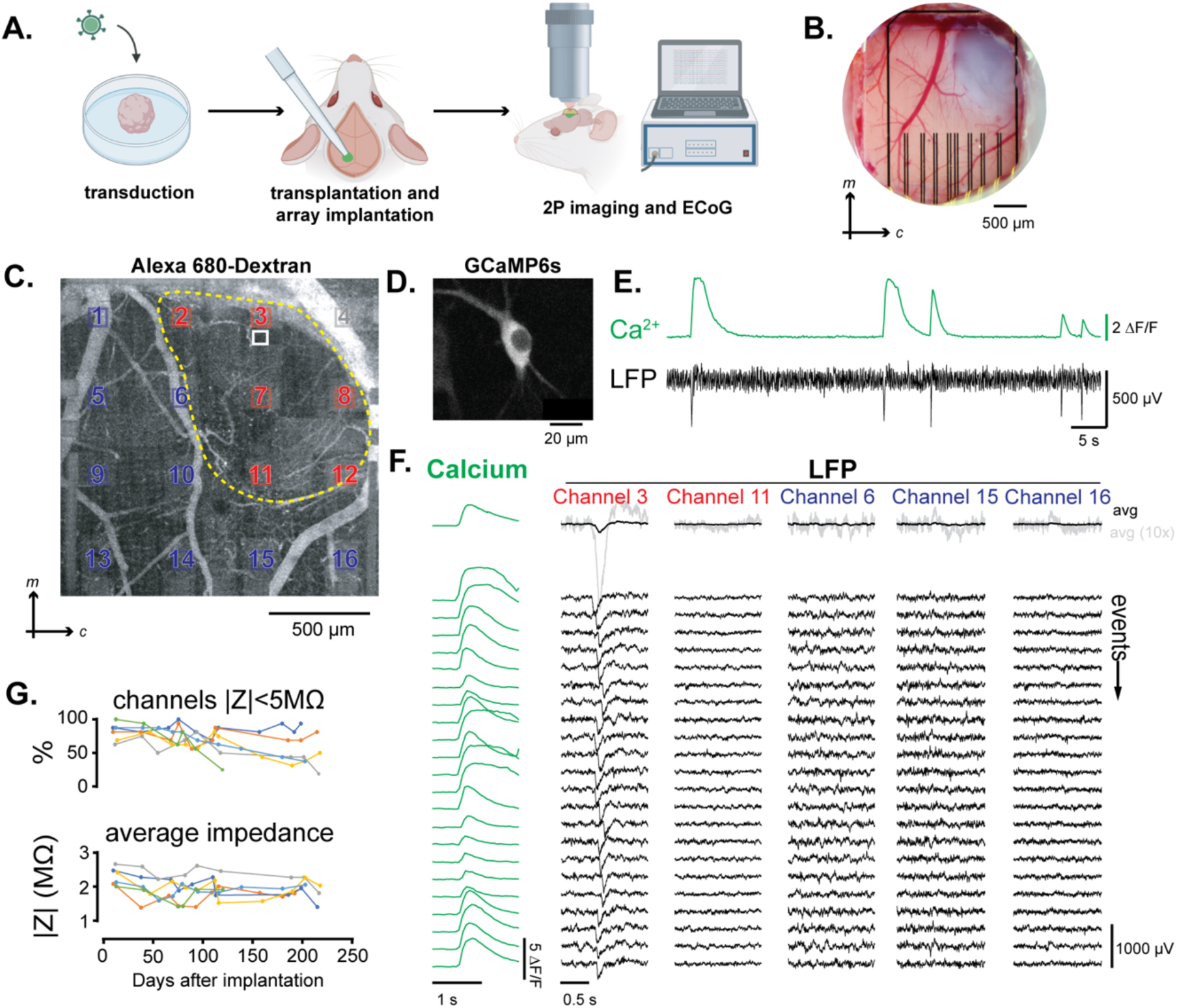
Local field potential (LFP) signals in electrocorticography (ECoG) recordings measured with graphene microelectrode arrays (gMEAs) coincide with calcium increases in human neurons and are spatially confined to the xenograft. A. Experimental paradigm for simultaneous two-photon (2P) calcium imaging and ECoG recordings. In this experiment, hCOs were transduced with AAV7m8 hSyn1-GCaMP6s-p2A-NLS-tdTomato in culture before xenotransplantation into retrosplenial cortex. Animals were recorded while awake and head-fixed without presentation of external stimuli. Schematic created in <u>BioRender</u>. B. Representative brightfield image taken at the end of the implantation surgery of the cranial exposure showing the implanted hCO xenograft and the gMEA. The gMEA is fixed to the glass window covering the exposure with optical-grade glue. Abbreviations: *m*, medial; *c*, caudal. C. Overview of the exposure acquired with 2P microscopy after labeling the blood plasma with Alexa 680-Dextran. Xenotransplant boundaries are indicated by the yellow dotted line. Graphene electrodes of the gMEA are highlighted in blue, red, and grey boxes that correspond to their location above host cortex, xenograft, and bone, respectively. The white box highlights the location of the calcium imaging field of view (FOV) for the data shown in panels D-F. D. GCaMP6s-expressing neuron recorded in the FOV shown in panel C. E. Excerpt of recorded calcium activity (shown as ΔF/F) of the neuron shown in panel D corresponding local field potential (LFP) signal recorded at the same time in channel 3 of the gMEA. F. (left) Calcium events detected in the neuron in panel D; events are aligned by calcium event onset. (right) Corresponding LFP signals in channels above the xenograft (channels 3 and 11) and above the host cortex (channels 6, 15, and 16). G. Proportion of graphene electrodes with an impedance <5 MΩ as a function of time in vivo (top) and average impedance of electrodes (with <5 MΩ) as a function of time in vivo (bottom); individual traces from six animals are shown.

## Discussion

Brain organoids represent an exciting avenue to study neurophysiology in a human-centric experimental context. Several recent studies have demonstrated that brain organoids xenotransplanted into the rodent cortex undergo vascularization, exhibit advanced neuronal maturation, and integrate into host neuronal circuits (Mansour *et al*. 2018, Revah *et al*. 2022, Wilson *et al*. 2022). However, most studies rely on a limited set of *in-vivo* tools and fail to fully capture the temporal trajectory of xenograft maturation across scales. Longitudinal functional and structural evaluation of these trajectories in the same xenograft is essential for faithfully studying normal brain development as well as the pathophysiology of neurodevelopmental and neuropsychiatric disorders. In the present study, we expand on previous work to establish a palette of optical tools for longitudinal assessments of xenograft structure and neuronal activity *in vivo*. Combining OCT with 2P, 1P, and SCAPE microscopy for calcium imaging over a 3-month period, we observed an at least 3-fold expansion in xenograft size, enhanced vascularization, functional maturation human neuronal activity patterns (e.g., network synchronization), and regional compartmentalization of large-scale calcium signals across the xenograft. Furthermore, simultaneous recording of calcium and electrical signals using gMEAs revealed spatial correlations between the activity of individual neurons and electrophysiological signals measured at the xenograft surface.

The proposed toolkit has several key advantages. Previously, xenograft vascularization has been measured via 2P microscopy using intravascular fluorescent dyes (Mansour *et al*. 2018, Shi *et al*. 2020, Wilson *et al*. 2022). However, early after xenograft implantation, dye leakage is more likely to occur due to increased blood-brain-barrier permeability of vessels entering the xenograft, as well as in the host vasculature in response to the surgery (Hawkins and Davis 2005, Hawkins and Egleton 2006), and intravascular injections increase the risk for a potentially fatal infection of the immunodeficient host animal. In this context, we established OCT as a label-free alternative for assessing xenograft vascularization earlier after implantation and at denser sampling intervals. OCT also achieves high contrast imaging of the host-graft interface which enables volumetric assessments of xenograft size, morphology, and positioning, representing a less expensive, easier alternative to MRI-based assessments, as shown, e.g., in Revah *et al*. (2022) and Wang *et al*. (2025). Additionally, while 2P microscopy has been used previously to assess single neurons in xenotransplanted hCOs via calcium imaging, here we use 1P widefield and SCAPE imaging to achieve mesoscale and volumetric calcium imaging, respectively, allowing us to assess neuronal activity in the hCO xenograft across scales and dimensions. Building on previous work (Mansour *et al*. 2018, Linaro *et al*. 2019, Revah *et al*. 2022, Wilson *et al*. 2022), we also demonstrate the feasibility of using gMEAs for host and graft ECoG recordings. Future work could leverage changes in gMEA size and geometry to enable ECoG recordings across a larger area of cortex surrounding the xenograft, or depth electrode arrays, such as the NeuroFITM design (Liu *et al*. 2021) could enable chronic recordings across xenograft depth and underlying structures, for example, the hippocampus.

Using our proposed toolkit, we observed neuronal activity that aligns with known neurodevelopmental milestones. Previous reports highlight that calcium signals in the cortex of fetal and newborn rodents transition from sparse and decorrelated to highly synchronous activity featuring longer calcium events across the cortex (Adelsberger *et al*. 2005, Wu *et al*. 2024). Synchrony at this stage is essential to promote activity-dependent refinement and synaptogenesis, and peaks in the rodent cortex at the time of birth (Corlew *et al*. 2004, Spitzer 2006, Pires *et al*. 2021). Subsequently, in postnatal stages a transition to desynchronized network activity takes place, leading to activity patterns like those in adult cortex (Golshani *et al*. 2009, Wu *et al*. 2024). In line with these reports, we observe synchronized calcium events, detectable by 2P and widefield 1P imaging, at approximately 3 months after transplantation. These results serve as a benchmark for neuronal network maturation of transplanted hCOs *in vivo* and can be used to illuminate perturbations of this model in future studies, e.g., in the context of a disease. Future studies should investigate extended time points to capture postnatal-like network transitions. Additionally, the spatial separation of signal components that we observed in widefield recordings may correspond to functionally distinct groups of neurons, potentially representing discrete functional ensembles and early cortical arealization (Jabaudon 2017). This functional compartmentalization aligns with previous observations of regional specialization in developing brain tissue (Warm *et al*. 2025) and suggests that widefield calcium imaging can effectively resolve such network structures in organoid xenografts. Lastly, the introduction of SCAPE microscopy allows us to study neuronal network connectivity in three-dimensional space, which is not possible using solely planar imaging modalities. This provides the opportunity to track spatiotemporal patterns in neuronal activity volumetrically across the xenograft. Future studies should leverage this to test the variation in neuronal activity patterns across depth and the possible emergence of laminar activity patterns.

Experiments in this study were performed using hCOs generated by two different protocols (Pasca *et al*. 2015, Trujillo *et al*. 2019), and a side-by-side comparison of organoids generated with different protocols and from different iPSC lines was beyond the scope of this study. While some findings on cell activity may not generalize, as they depend on organoid type and cell origin, this does not affect the usability or performance of the methods we describe herein. Variability across organoids and consequently across xenografts has been recognized as a potential shortcoming, limiting the interpretability and translational relevance of xenotransplantation models. *In vivo* tracking methods following graft integration and maturation in a chronic preparation, as presented here, could help guide the development of technologies, such as synthetic gene regulation, optogenetics, microfluidics, and biomaterials, that aim to improve reproducibility across samples.

Here, we primarily track changes in xenograft structure and activity, but do not follow corresponding changes in the host brain. This could be enabled, for example, via two-color calcium imaging in host and xenograft with spectrally compatible indicators. While a large variety of transgenic mice with respective reporter transgenes is available (Arias *et al*. 2022), intercrossing the immunodeficient NOD/SCID line with reporter lines typically maintained on a C57Bl/6 background is a considerable effort. Alternative strategies include local AAV delivery into the host brain or systemic delivery of AAVs capable of crossing the blood-brain barrier (Chen *et al*. 2022) into NOD/SCID mice, as recently reported by Drexler *et al*. (2025).

The toolkit we propose here provides an opportunity to longitudinally investigate other xenograft systems *in vivo* in a chronic preparation. Recent advances in brain organoid technology have included the incorporation of additional cell types, such as mural cells and microglia (Fagerlund *et al*. 2022, Zhang *et al*. 2023), and the emergence of assembloids generated via fusion of regionally-specified organoids (e.g., hCOs with dorsal and ventral identities) (Birey *et al*. 2017). These systems offer new opportunities to investigate mechanisms of neurodevelopment, such as the role of microglia in shaping neuronal circuitry (Kettenmann *et al*. 2011) or migration of GABAergic interneurons within the developing cortex, as well as their role in neuronal network state transitions during cortical development (Wonders and Anderson 2006). Xenotransplantation of these advanced organoid systems into the rodent brain has recently been demonstrated (Schafer *et al*. 2023, Wang *et al*.). Future studies should employ strategies for longitudinal, *in-vivo* assessments of these systems, such as the toolkit proposed herein, to wholistically investigate their temporal maturation.

Taken together, our neurophotonics toolkit represents a substantial advancement in our ability to comprehensively assess structural and functional changes in organoid xenografts. We predict that our platform will enable further studies using organoid xenotransplantation to investigate human neurodevelopmental pathologies such as autism spectrum disorder, epilepsy, and Down syndrome; screen therapeutic interventions; and monitor organoid-mediated tissue restoration. Insights gained from these studies promise to advance our understanding of neurodevelopmental pathologies, inform treatment strategies, and improve clinical outcomes.

## Methods

### hCO culture and viral transduction

For this study, hCOs were generated and cultured using one of two previously established protocols: Protocol A (Trujillo *et al*. 2019, Fitzgerald *et al*. 2024) and Protocol B (Pasca *et al*. 2015). Cell lines and protocols used for each xenotransplantation experiment are listed in **Supplementary Table 1.** For fluorescence imaging, hCOs were transduced using either LV or AAV vectors (**Supplementary Table 1**). Information for each virus and titers used can be found below in **Table 1**.

**Table 1.** Viral vectors used in this study.

| Construct | Titer | Product Information |
| --- | --- | --- |
| AAV7m8-hSyn1-GCaMP6s-P2A-nls-dTomato | $8.4 \times 10^{12}$ vg/mL | Addgene, 51084; AAV7m8 produced by Viral Vector Production Unit, UAB Centre de Biotechnologia Animal i de Teràpia Gènica, Universitat Autònoma Barcelona. |
| pGP-AAV1-synjGCaMP8s-WPRE | $\geq 1 \times 10^{13}$ vg/mL | Addgene, 162374-AAV1 |
| pAAV8-CAG-tdTomato (codon diversified) | $\geq 1 \times 10^{13}$ vg/mL | Addgene, 59462-AAV8 |
| pLV[Exp]-Puro-SYN1.jGCaMP8s | $\geq 1 \times 10^9$ TU/mL | VectorBuilder, VB240904-1449tnm |
| pLV[Exp]-Bsd-EF1A>mScarlet3 | $\geq 1 \times 10^9$ TU/mL | VectorBuilder, VB240919-1466xpw |

For hCOs generated by protocol A, viral transduction was performed at mid- to late-stages of differentiation to ensure robust neuronal maturation, as previously described (Fitzgerald *et al*. 2024). Briefly, healthy hCO (∼40–45 DIV) with evident neural rosettes were selected and pooled (≈20 organoids per well) and exposed to AAV in suspension. hCOs were incubated with AAV7m8 vectors encoding genetically encoded calcium indicators (AAV-hSyn1-GCaMP6s-P2A-nls-dTomato) at a final dose of ∼1×10¹⁰ viral genomes per well (≈1×10⁹ vg per organoid) in fresh M2 maintenance medium (Neurobasal with 1% (v/v) GlutaMAX, 1% (v/v) MEM non-essential amino acids, and 2% (v/v) Gem21). Following a 5–6 h incubation at 37 °C on an orbital shaker, conditioned M2 medium was added to support recovery, and organoids were returned to standard culture conditions with media changes every 3-4 days. Starting on 45 DIV, hCOs with robust and stable transgene expression were used for transplantations.

For hCOs generated by Protocol B, hCOs at 25-30 DIV were transduced as previously described (Kelley *et al*. 2024). For viral infections, hCOs were moved to a 1.5–mL tube and incubated for 30 min at 37 °C in 20 μL of maintenance media with 0.5 μL or 4 μL of AAV or LV, respectively (**Table 1**). After 30 min of incubation, 300 μL of maintenance media was added, and the hCOs were incubated at 37 °C overnight. On the following morning, hCOs were moved back to 24-well plates until the grafting procedure. Starting on 40 DIV, verified fluorescent hCOs were used for transplantations.

### Experimental animals

All experiments and animal procedures were conducted according to the Guide for the Care and Use of Laboratory Animals and protocols approved by the Boston University Institutional Animal Care and Use Committee (PROTO202000026). We used non-obese diabetic, severe combined immunodeficient (NOD/SCID) mice (Charles River Laboratories) at an age of 12-20 weeks. Matching the sex of the xenografted hCOs, we used female mice in this study. Animals were single- or group-housed in isolated ventilated rodent cages under normal light-dark cycles with nestlet enrichment and unrestricted access to food and water.

We performed a single 90-120-min long surgery for headbar placement, craniotomy and durectomy, aspiration of retrosplenial cortex, organoid implantation, chronic gMEA implantation, and closure of the exposure with a glass window. Animals received 4.8 mg/kg Dexamethasone, 1 mg/kg slow-release Buprenorphine, and 5 mg/kg Meloxicam 120-180 min before surgery. Animals were anesthetized using 90 mg/kg Ketamine and 10 mg/kg Xylazine, or a cocktail of 0.05 mg/kg Fentanyl, 0.25 mg/kg Dexmedetomidine, and 5 mg/kg Midazolam. For the duration of the surgery, animals were placed within a stereotaxic frame on a feedback-controlled heating blanket and received 100% oxygen through a nose cone. After preparation of the animal, the skin overlying the dorsal cranium was removed, and wound borders were closed with surgical-grade cyanoacrylate glue (VetBond Tissue Adhesive, 3M). The skull was cleaned and etched with phosphoric acid gel for 60-90 s and cleaned with 0.9% NaCl in water. Then, a thin layer of UV-curable primer (OptiBond, Kerr Dental) was applied. After fixing the custom-made titanium headbar to the bone using UV-curable dental resin (Tetric EvoFlow, Ivoclar), a 4-mm large piece of bone above the left retrosplenial cortex and the underlying dura mater were removed. Then, a cylindrical piece of retrosplenial cortex (diameter: ca. 1-1.5 mm, length: 1 mm) was removed via aspiration with a blunt needle. After controlling bleeding using hemostatic sponges, we placed a single organoid inside the cavity. If needed, the organoid was cut in half using micro-scissors, and one fragment of the organoid was implanted. Then, we used a glass window to which a transparent graphene electrode array was optionally fixed using UV-curable glue (NOA 61, Norland) to cover the exposure. The window was held in place using a stereotaxic arm, and after removal of excess fluid, dental resin (Tetric EvoFlow, Ivoclar) was used to seal the exposure and fix the glass window to the skull. A removable, custom 3D-printed protective cap was fixed to the headbar to protect exposure and microelectrode array from damage (Wilson *et al*. 2022). Immediately after surgery, animals received a single dose of 500 mg/kg Cefazolin; Meloxicam (5 mg/kg every 24 h) was administered for three days after surgery. Animals received Sulfamethoxazole/Trimethoprim (Sulfatrim) either via drinking water (0.53 mg/mL Sulfamethoxazole and 0.11 mg/mL Trimethoprim) for three days before and up-to 6 weeks after surgery, or via medicated food (Uniprim diet, Envigo, TD06596) for the duration of the study.

After a recovery period of seven days, animals underwent habituation in one session per day to accept increasingly longer periods (up-to 120 min) of head restraint inside the microscope enclosure. Drops of sugar water (infusion-grade 5% dextrose in 0.9% NaCl) were offered as a reward during training and recording sessions.

### Two-photon imaging of vessel structure and calcium dynamics

#### Data acquisition

Two-photon imaging was performed in awake, head-fixed animals on commercial two-photon laser scanning microscope systems (Bruker Ultima, Bruker Ultima Investigator Plus) with Coherent Chameleon Ultra II or Coherent Chameleon Discovery Ti:Sapphire lasers tuned to 920-950 nm for excitation of GCaMP6s or 8s. For vascular imaging, in-house produced Alexa 680-Dextran (using amino-dextran with 0.5 MDa molecular weight, Fina Biosolutions) was injected under isoflurane anesthesia 15-30 min before imaging. Overview images of the cranial window were acquired with a 4× objective (Plan-Neofluar, NA = 0.16, Zeiss) at the beginning of each imaging session. For high-resolution and functional imaging, a 20× objective (XLUMPlanFLNXW, NA = 1.0, Olympus) was used. Calcium imaging was performed in consecutive 180-300-s long acquisition runs either with two conventional galvo mirrors in custom-defined regions of interest of variable size at imaging rates of 5-15 Hz, or in galvo/resonant galvo mode with an image size of 512 × 512 pixels and an acquisition rate of 30 Hz; the average of two consecutive frames was is stored in the final image series, resulting in an effective frame rate of 15 Hz.

#### Data postprocessing

Image time series were stored as Tiff files and then imported together with the corresponding metadata into MATLAB. Imaging time series underwent in-plane motion correction with ‘normcorre’ (Pnevmatikakis and Giovannucci 2017); in most cases, images from a reference channel showing vasculature or fluorescence of a ‘static’ fluorophore were used; estimated shifts were then applied to all channels. Regions of interest were either manually delineated in MATLAB, or time series were exported into Suite2P (Pachitariu *et al*. 2017) for automatic ROI identification with Cellpose using the setting ‘anatomical_only=2’. For image series acquired in resonance scanning mode, rapid, non-physiological changes in GCaMP signal intensity due to tissue motion along the Z axis were identified using a custom-written MATLAB GUI. Here, manual thresholding of time courses - and their temporal derivatives – from the brightest 5% of pixels (or regions of interest) from a reference channel (i.e., vasculature or structural fluorophores) was used to identify frames affected by motion. After identifying and flagging motion-affected frames, time series were down-sampled to 2 Hz for further analysis. For each recording, fluorescence changes in neuronal cell bodies were extracted as ΔF/F. Each trace was binarized into periods of activity (calcium events) or inactivity based on a threshold of ten standard deviations of the signal baseline.

#### Calcium imaging quantification

Quantification of 2P calcium imaging data was performed on binarized traces using custom scripts in MATLAB 2024b. Extracted metrics for imaging trial included fraction of active cells, average active time, event frequency, synchronicity, event length, event height, and the CV for the inter-event interval, IEI. Active cells were defined as those having at least one calcium event (defined above). Synchronicity was measured using Pearson’s correlation coefficient across cells within the same field of view.

### Optical coherence tomography imaging and quantitative analysis

#### Data acquisition

For OCT imaging, a spectral domain OCT system was used (1310 nm center wavelength, 170 nm bandwidth, Thorlabs). OCT imaging was performed as described previously (Adewumi *et al*. 2025). In brief, images were acquired for the entire optical window (3 mm × 3 mm) using a 5× objective (NA = 0.14, Mitutoyo) with scans of 1000 × 1000 pixels. B-scans were repeated twice and averaged or subtracted, then rastered into C-scans. C-scans were repeated twice and averaged to generate intensity profiles or angiograms respectively. Images were log-transformed for improved contrast.

#### Angiogram quantification

For quantification of capillary density, ROIs (1 mm × 1 mm) were imposed onto 500-µm MIPz angiograms within the graft boundary. ROIs were then analyzed for total capillary network length using the open-source application Angiotool (Zudaire *et al*. 2011). The reported total capillary network length for each ROI was divided by the ROI area to determine the network density.

#### Intensity profile quantification

For quantification of xenograft volume, MIPz images were generated for the entire intensity profile image in 50-µm segments. For each MIPz, an ROI was drawn around the graft boundary. ROIs were then stacked and interpolated to create one continuous 3D object and gaussian smoothing was applied (σ=10). A volume was approximated based on pixel size, and data was visualized using MATLAB (MATLAB 2024b).

### Widefield mesoscopic imaging

Widefield imaging was performed in awake, head-fixed animals on a custom-built four-channel widefield microscope as previously described (Doran *et al*. 2024) in acquisition runs of 300-1200 s with minor modifications: the acquisition frequency for one four-channel cycle was 15 Hz and images were binned on the camera chip by a factor of 2×2, corresponding to a spatial resolution of 10 µm per pixel.

Raw image data was imported into MATLAB (2024b) and split into four individual image time series (green fluorescence, F_470_, red fluorescence, F_565_, reflection at 530 nm, R_530_, and reflection at 625 nm, R_625_). Detection of in-plane motion was performed in MATLAB using ‘normcorre’ (Pnevmatikakis and Giovannucci 2017) using the R_530_ time-series after applying a spatial high-pass filter; estimated image shifts were then applied to all other time series. For further analysis, changes in (dynamic) green fluorescence (F_470_) were normalized to (static) red fluorescence (F_565_) as F*_470_=F^N^_470_(x,y,)-f(x,y)*F^N^_565_(x,y) with f(x,y)=*mldivide*(F^N^_565_(x,y), F^N^_470_(x,y)), where F^N^_470_=F_470_-avg_t_(F_470_), F^N^_565_=F_565_-avg_t_(F_565_), with avg_t_ as temporal average and x, y as pixel coordinates.

For event interval analysis, we used F*_470_ signal intensities averaged across all pixels from a region of interest manually defined from F_470_ and F_565_ images. Before peak detection using MATLAB’s ‘findpeaks’ function, low-frequency drifts in baseline fluorescence, e.g., due to photobleaching, were corrected via baseline subtraction. For principal component analysis (PCA) using MATLAB’s ‘*pca*’ function, a manually drawn mask outlining the entire exposure was defined, and after spatial down-sampling the images with a factor of four, baseline drifts were corrected via pixel-wise subtraction of a baseline time course calculated from the exposure-wide average of the F*_470_ signal.

### Swept confocally-aligned planar excitation (SCAPE) imaging

SCAPE imaging was performed in awake, head-fixed animals using a custom-built SCAPE microscope, based on the design of the Hillman Laboratory (Voleti *et al*. 2019). Recordings were made using a 20× water immersion objective (XLUMPlanFLNXW, NA = 1, Olympus). GCaMP8s fluorescence was excited at 488 nm. Volumetric datasets were acquired at ten volumes per second (VPS) with a field of view of 395 µm × 340 µm × 171 µm.

Raw data were reconstructed and de-skewed using a custom MATLAB pipeline previously developed by the Hillman Laboratory. Subsequent analysis was performed using a custom Python-based imaging toolbox for 3D time-series data processing. The pipeline includes ΔF/F normalization, ROI segmentation, and trace extraction. The baseline fluorescence for each voxel was defined as the 20th percentile of its intensity distribution over time. Three-dimensional regions of interest (ROIs) corresponding to neuronal cell bodies were identified through intensity-based thresholding, morphological filtering, and event-linked segmentation, followed by semi-manual quality control and visualization in Napari (Chiu and Clack 2022).

### Graphene Microelectrode Array Fabrication

Graphene microelectrode arrays were fabricated following our previous fabrication protocols (Liu *et al*. 2017, Thunemann *et al*. 2018, Wilson *et al*. 2022) with an added second layer of graphene and nitric acid doping to lower electrode impedance (Ramezani *et al*. 2024). To start, polydimethylsiloxane (PDMS) was spun onto four-inch silicon wafers and annealed on a 150 °C hotplate for 10 min. A 50-µm-thick sheet of PET was then placed on the adhesive PDMS layer as the electrode array substrate. Cr/Au wires (10- and 100-nm-thick, respectively) were deposited onto the PET using a Denton Discovery 18 Sputtering System. The metal wires were then patterned using photolithography (Heidelberg MLA150) and wet etching (Gold Etchant TFA, Chromium Etchant 1020AC). Monolayer graphene was transferred onto the wafer using an electrochemical delamination process (Wang *et al*. 2011). The sacrificial transfer substrate, poly(methyl methacrylate), was removed with acetone and IPA and the wafer was submerged in 50% HNO_3_ solution for 10 minutes to dope the graphene and lower its sheet resistance. A second monolayer was transferred onto the wafer and cleaned using acetone and isopropyl alcohol (IPA). Graphene wires were patterned using PMGI/AZ1512 bilayer photolithography and oxygen plasma etching (Plasma Etch PE100). SU-8 2005 was spun onto the wafer as an encapsulation layer and openings were patterned using photolithography. Finally, the wafer was cleaned using an alternating ten-minute IPA/DI water rinse then baked in ramped intervals from 125 to 135 °C to seal the SU-8 encapsulation layer. The PET substrate was peeled from the PDMS-coated wafer and arrays were cut out for characterization and animal experiments. EIS characterizations were conducted in 1x PBS using the Gamry Reference 600+ system. Additionally, arrays were deposited with platinum nanoparticles to further lower impedance to 600 kΩ by placing them in a H_2_PtCl_6_ (0.05 M) and K_2_HPO_4_ (0.01 M) solution in a two-electrode configuration and flowing 50 nA of current between electrodes and a counter electrode for eight seconds.

### Electrophysiological recordings

After head fixation and removing the protective cap, the array was connected to a flexible ribbon connector to a PCB equipped with a dual row horizontal Nano Strip connector (Omnetics Connector Corporation) for connection to the recording headstage (RHD 32-channel Recording Headstage, Intan Technologies) and ground and reference inputs. Miniature alligator clips were used to connect the reference screw in the mouse skull to the PCB board. The ground input on the PCB was connected to a screw of the head fixation system. Recordings were performed with a RHD Recording Controller or Interface Board (Intan Technologies) with a sampling rate of 20 kHz. For synchronization with external instrumentation such as behavioral camera or two-photon microscope during data processing, timing triggers generated by a dedicated digital-analog interface or the microscope system were recorded as analog or digital signals by the electrophysiology recording system.

Raw electrophysiology data was imported and further analyzed in MATLAB. Independent component analysis (ICA) was performed using the *jadeR* function adapted from the publicly available MATLAB-based EEGLab resource (https://eeglab.org/) (Delorme and Makeig 2004) to mitigate movement-related artifacts in the electrophysiological data. We removed up to three independent components that were (1) lacking spatial heterogeneity and (2) followed the time-course of body movement or the scanning pattern of the microscope system. Raw electrophysiological data were low pass filtered at 250 Hz and resampled to 4000 Hz to isolate the local field potential (LFP).

### Monitoring and recording of mouse behavior

A CCD camera (acA1920-150um, Basler) attached to a variable zoom lens (7000 Macro Lens, Navitar) and 940-nm LED (M940L3, Thorlabs) were used to monitor behavior and motion movement of awake head-fixed mice during electrophysiology, single-photon- and two-photon imaging. The CCD camera was externally triggered by a dedicated digital-analog interface or the microscope system to synchronize recordings of mouse behavior and other modalities.

In addition, an accelerometer (ADXL335, Analog Devices) was placed underneath the mouse body. Analog signals from the accelerometer were recorded either through a dedicated digital-analog interface or the electrophysiology recording system.

### Immunohistochemistry (IHC)

Animals were sacrificed by barbiturate overdose and immediately transcardially perfused using 0.2% heparin in phosphate-buffered saline (PBS) for 4 minutes followed by 4% paraformaldehyde (PFA) for 4 minutes at a rate of 5 mL/min. Brains were harvested and kept at 4 °C in 4% PFA for 12 hours. Brains were then cryoprotected in 1x TRIS-buffered saline (TBS, pH=7.4) with 30% sucrose and 0.001% sodium azide for 4 days at 4 °C. Brains were embedded in optimal cutting temperature (OCT) compound and sectioned coronally (30-40 µm) using a cryostat (CM3050, Leica). Sliced sections were stored free floating in 1x TBS with 0.001% sodium azide at 4 °C.

Immunofluorescent staining was conducted using a free-floating staining protocol (Adewumi *et al*. 2025). Tissue was treated with 1N hydrochloric acid for 10 minutes for antigen retrieval, then washed with 1x TBS. Then, sections were blocked and permeabilized and 5-10% normal donkey serum and 0.5% Triton X-100 in 1x TBS for one hour. Brain sections were incubated in primary antibody solution overnight at 4 °C. The primary antibodies used in experiments are chicken anti-GFP (1:500, cat# ab13970, Abcam); rabbit anti-HNA (1:250, cat# NBP3-13912, Novus Biologicals); mouse anti-NeuN (1:200, cat# MAB377, Sigma-Aldrich), mouse anti-STEM121 (1:500, cat #Y40410, Takara Bio). Tissue was again washed and incubated in the secondary antibody solution containing 5-10% donkey serum for 2 h at room temperature. Secondary antibodies (FITC, TRITC, Cy5, and Cy7) were sourced from Jackson ImmunoResearch Laboratories with a donkey host animal, and target as specified by the corresponding primary antibody. After additional washing, DAPI staining was performed for 10 minutes (DAPI; 2 ng/mL; Molecular Probes). Brain sections were mounted onto slides using ProLong Gold anti-fade reagent (Molecular Probes). Stained sections were imaged using a spinning disk confocal microscope (IX83, Olympus).

### Quantification of IHC data

Cell detection was performed using the “Cell detection” function in QuPath (v0.5.1). Cell counts were calculated for manually drawn ROIs around the boundary of the xenograft based on the given marker (HNA, NeuN or GFP). NeuN+ and GFP+ cells were counted when colocalized with HNA. NeuN+ and GFP+ cells were normalized to HNA+ cell numbers.

### Statistical analysis

GraphPad Prism (v10.4.2) was used to perform Student’s t-tests with significance being defined as p-values of less than 0.05. Analysis of longitudinal trends in calcium imaging data was conducted using linear mixed effects models (LMM), and principal component analysis (PCA) was performed where indicated. Both LMM and PCA analyses were performed using MATLAB 2024b.

## Resource Availability

Additional data and scripts used for data analysis are available upon request.

## Acknowledgements

We thank members of the Devor/Thunemann, Zeldich, and O’Shea labs for insightful discussions and technical support, Dr. Rebeca Blanch (UCSD) for help with the AAV7m8 transduction protocol, Dr. Omer Revah (Hebrew University of Jerusalem) for his invaluable advice regarding *in vitro* calcium imaging and viral labelling strategies, Dr. Steven J. Haggarty (Harvard Medical School and Massachusetts General Hospital) for kindly providing the MGH2046 hIPSC line, and the Boston University Neurophotonics Center for access to imaging facilities and technical support. This work was supported by grants from the National Institutes of Health (NIH) including: RF1AG088529 (to E.Z. and M.T.); R21AG080269 (to E.Z.); R01MH107367, R01HD107788, R01NS105969, and R01NS123642 (to A.R.M). K.E.H. is supported by the NIH F31 Fellowship 1F31NS141509-01A1, the NIH Quantitative Biology & Physiology training program T32GM145455, and the Boston University Neurophotonics Center CAN DO award. Other support includes funding from the California Institute for Regenerative Medicine DISC2-13515 (to A.R.M.) and from the Department of Defense W81XWH2110306 (to A.R.M.).

## Author Contributions

**Kate E. Herrema**: Conceptualization, Investigation, Formal Analysis, Data Curation, Writing - Original Draft, Writing - Review & Editing; **Elizabeth K. Kharitonova**: Conceptualization, Investigation, Formal Analysis, Data Curation, Writing - Original Draft, Writing - Review & Editing; **Daria Bogatova**: Investigation, Formal Analysis, Software, Writing - Review & Editing; **Madison Wilson**: Investigation, Resources, Writing - Review & Editing; **Emily A. Martin**: Investigation, Writing - Review & Editing; **Clara Chung**: Investigation, Writing - Review & Editing; **Francesca Puppo**: Investigation, Resources, Writing - Review & Editing; **Shira Klorfeld-Auslender**: Resources, Writing - Review & Editing; **Pierre Boucher**: Software, Formal Analysis, Writing - Review & Editing; **Li-Huei Tsai**: Resources, Supervision, Writing - Review & Editing; **Alysson R. Muotri**: Resources, Methodology, Funding acquisition, Supervision, Writing - Review & Editing; **Duygu Kuzum**: Conceptualization, Funding acquisition, Methodology, Resources, Supervision, Writing - Review & Editing; **Anna Devor**: Conceptualization, Funding acquisition, Resources, Supervision, Writing - Review & Editing; **Timothy O’Shea**: Conceptualization, Funding acquisition, Methodology, Resources, Supervision, Writing - Original Draft, Writing - Review & Editing; **Ella Zeldich**: Conceptualization, Funding acquisition, Methodology, Resources, Supervision, Writing - Original Draft, Writing - Review & Editing; **Martin Thunemann**: Conceptualization, Formal Analysis, Investigation, Methodology, Software, Supervision, Writing - Original Draft, Writing - Review & Editing.

## Declaration of interests

Dr. Muotri is a co-founder and has an equity interest in TISMOO, a company dedicated to genetic analysis and human brain organogenesis, focusing on therapeutic applications customized for the autism spectrum disorder and other neurological conditions. Dr. Muotri is also an inventor on several patents using brain organoids. The terms of this arrangement have been reviewed and approved by the University of California, San Diego, in accordance with its conflict-of-interest policies.

## References

Adelsberger, H., O. Garaschuk and A. Konnerth (2005). “Cortical calcium waves in resting newborn mice.” Nature Neuroscience 8(8): 988–90. [10.1038/nn1502]

Adewumi, H. O., M. G. Simkulet, G. Kureli, J. T. Giblin, A. B. Lopez, S. E. Erdener, J. Jiang, D. A. Boas and T. M. O’Shea (2025). “Optical coherence tomography enables longitudinal evaluation of cell graft-directed remodeling in stroke lesions.” Exp Neurol 385: 115117. [https://www.ncbi.nlm.nih.gov/pubmed/39694221]

Amiri, A., G. Coppola, S. Scuderi, F. Wu, T. Roychowdhury, F. Liu, S. Pochareddy, Y. Shin, A. Safi, L. Song, Y. Zhu, A. M. M. Sousa, E. C. Psych, M. Gerstein, G. E. Crawford, N. Sestan, A. Abyzov and F. M. Vaccarino (2018). “Transcriptome and epigenome landscape of human cortical development modeled in organoids.” Science 362(6420): eaat6720. [https://www.ncbi.nlm.nih.gov/pubmed/30545853]

Arias, A., L. Manubens-Gil and M. Dierssen (2022). “Fluorescent transgenic mouse models for whole-brain imaging in health and disease.” Front Mol Neurosci 15: 958222. [https://www.ncbi.nlm.nih.gov/pubmed/36211979]

Birey, F., J. Andersen, C. D. Makinson, S. Islam, W. Wei, N. Huber, H. C. Fan, K. R. C. Metzler, G. Panagiotakos, N. Thom, N. A. O’Rourke, L. M. Steinmetz, J. A. Bernstein, J. Hallmayer, J. R. Huguenard and S. P. Pasca (2017). “Assembly of functionally integrated human forebrain spheroids.” Nature 545(7652): 54–9. [https://www.ncbi.nlm.nih.gov/pubmed/28445465]

Bouchard, M. B., V. Voleti, C. S. Mendes, C. Lacefield, W. B. Grueber, R. S. Mann, R. M. Bruno and E. M. Hillman (2015). “Swept confocally-aligned planar excitation (SCAPE) microscopy for high speed volumetric imaging of behaving organisms.” Nat Photonics 9(2): 113–9. [https://www.ncbi.nlm.nih.gov/pubmed/25663846]

Camp, J. G., F. Badsha, M. Florio, S. Kanton, T. Gerber, M. Wilsch-Brauninger, E. Lewitus, A. Sykes, W. Hevers, M. Lancaster, J. A. Knoblich, R. Lachmann, S. Paabo, W. B. Huttner and B. Treutlein (2015). “Human cerebral organoids recapitulate gene expression programs of fetal neocortex development.” Proc Natl Acad Sci U S A 112(51): 15672–7. [https://www.ncbi.nlm.nih.gov/pubmed/26644564]

Cardin, J. A., M. C. Crair and M. J. Higley (2020). “Mesoscopic Imaging: Shining a Wide Light on Large-Scale Neural Dynamics.” Neuron 108(1): 33–43. [https://www.ncbi.nlm.nih.gov/pubmed/33058764]

Chen, X., S. Ravindra Kumar, C. D. Adams, D. Yang, T. Wang, D. A. Wolfe, C. M. Arokiaraj, V. Ngo, L. J. Campos, J. A. Griffiths, T. Ichiki, S. K. Mazmanian, P. B. Osborne, J. R. Keast, C. T. Miller, A. S. Fox, I. M. Chiu and V. Gradinaru (2022). “Engineered AAVs for non-invasive gene delivery to rodent and non-human primate nervous systems.” Neuron 110(14): 2242–57 e6. [https://www.ncbi.nlm.nih.gov/pubmed/35643078]

Chiu, C.-L. and N. Clack (2022). “napari: a Python Multi-Dimensional Image Viewer Platform for the Research Community.” Microscopy and Microanalysis 28(S1): 1576–7. [10.1017/S1431927622006328]

Corlew, R., M. M. Bosma and W. J. Moody (2004). “Spontaneous, synchronous electrical activity in neonatal mouse cortical neurones.” J Physiol 560(Pt 2): 377–90. [https://www.ncbi.nlm.nih.gov/pubmed/15297578]

Daviaud, N., R. H. Friedel and H. Zou (2018). “Vascularization and Engraftment of Transplanted Human Cerebral Organoids in Mouse Cortex.” eNeuro 5(6): ENEURO.0219–18.2018. [https://www.ncbi.nlm.nih.gov/pubmed/30460331]

Delorme, A. and S. Makeig (2004). “EEGLAB: an open source toolbox for analysis of single-trial EEG dynamics including independent component analysis.” J Neurosci Methods 134(1): 9–21. [https://www.ncbi.nlm.nih.gov/pubmed/15102499]

Doran, P. R., N. Fomin-Thunemann, R. P. Tang, D. Balog, B. Zimmerman, K. Kilic, E. A. Martin, S. Kura, H. P. Fisher, G. Chabbott, J. Herbert, B. C. Rauscher, J. X. Jiang, S. Sakadzic, D. A. Boas, A. Devor, I. A. Chen and M. Thunemann (2024). “Widefield in vivo imaging system with two fluorescence and two reflectance channels, a single sCMOS detector, and shielded illumination.” Neurophotonics 11(3): 034310. [https://www.ncbi.nlm.nih.gov/pubmed/38881627]

Drexler, R., A. Drinnenberg, A. Gavish, B. Yalcin, K. Shamardani, A. E. Rogers, R. Mancusi, V. Trivedi, K. R. Taylor, Y. S. Kim, P. J. Woo, N. Soni, M. Su, A. Ravel, E. Tatlock, A. Midler, S. H. Wu, C. Ramakrishnan, R. Chen, A. E. Ayala-Sarmiento, D. R. Fernandez Pacheco, L. Siverts, T. L. Daigle, B. Tasic, H. Zeng, J. J. Breunig, K. Deisseroth and M. Monje (2025). “Cholinergic neuronal activity promotes diffuse midline glioma growth through muscarinic signaling.” Cell 188(17): 4640–57 e30. [https://www.ncbi.nlm.nih.gov/pubmed/40541184]

Fagerlund, I., A. Dougalis, A. Shakirzyanova, M. Gómez-Budia, A. Pelkonen, H. Konttinen, S. Ohtonen, M. F. Fazaludeen, M. Koskuvi, J. Kuusisto, D. Hernández, A. Pebay, J. Koistinaho, T. Rauramaa, Š. Lehtonen, P. Korhonen and T. Malm (2022) “Microglia-like Cells Promote Neuronal Functions in Cerebral Organoids.” Cells 11, 124 DOI: 10.3390/cells11010124.

Fischer, J., M. Heide and W. B. Huttner (2019). “Genetic Modification of Brain Organoids.” Front Cell Neurosci 13: 558. [https://www.ncbi.nlm.nih.gov/pubmed/31920558]

Fitzgerald, M. Q., T. Chu, F. Puppo, R. Blanch, M. Chillon, S. Subramaniam and A. R. Muotri (2024). “Generation of ‘semi-guided’ cortical organoids with complex neural oscillations.” Nat Protoc 19(9): 2712–38. [https://www.ncbi.nlm.nih.gov/pubmed/38702386]

Golshani, P., J. T. Goncalves, S. Khoshkhoo, R. Mostany, S. Smirnakis and C. Portera-Cailliau (2009). “Internally mediated developmental desynchronization of neocortical network activity.” J Neurosci 29(35): 10890–9. [https://www.ncbi.nlm.nih.gov/pubmed/19726647]

Gordon, A., S. J. Yoon, S. S. Tran, C. D. Makinson, J. Y. Park, J. Andersen, A. M. Valencia, S. Horvath, X. Xiao, J. R. Huguenard, S. P. Pasca and D. H. Geschwind (2021). “Long-term maturation of human cortical organoids matches key early postnatal transitions.” Nat Neurosci 24(3): 331–42. [https://www.ncbi.nlm.nih.gov/pubmed/33619405]

Hawkins, B. T. and T. P. Davis (2005). “The blood-brain barrier/neurovascular unit in health and disease.” Pharmacol Rev 57(2): 173–85. [https://www.ncbi.nlm.nih.gov/pubmed/15914466]

Hawkins, B. T. and R. D. Egleton (2006). “Fluorescence imaging of blood-brain barrier disruption.” J Neurosci Methods 151(2): 262–7. [https://www.ncbi.nlm.nih.gov/pubmed/16181683]

Jabaudon, D. (2017). “Fate and freedom in developing neocortical circuits.” Nat Commun 8(1): 16042. [https://www.ncbi.nlm.nih.gov/pubmed/28671189]

Jgamadze, D., J. T. Lim, Z. Zhang, P. M. Harary, J. Germi, K. Mensah-Brown, C. D. Adam, E. Mirzakhalili, S. Singh, J. B. Gu, R. Blue, M. Dedhia, M. Fu, F. Jacob, X. Qian, K. Gagnon, M. Sergison, O. Fruchet, I. Rahaman, H. Wang, F. Xu, R. Xiao, D. Contreras, J. A. Wolf, H. Song, G. L. Ming and H. I. Chen (2023). “Structural and functional integration of human forebrain organoids with the injured adult rat visual system.” Cell Stem Cell 30(2): 137–52 e7. [https://www.ncbi.nlm.nih.gov/pubmed/36736289]

Kadoshima, T., H. Sakaguchi, T. Nakano, M. Soen, S. Ando, M. Eiraku and Y. Sasai (2013). “Self-organization of axial polarity, inside-out layer pattern, and species-specific progenitor dynamics in human ES cell-derived neocortex.” Proc Natl Acad Sci U S A 110(50): 20284–9. [https://www.ncbi.nlm.nih.gov/pubmed/24277810]

Kelley, K. W., O. Revah, F. Gore, K. Kaganovsky, X. Chen, K. Deisseroth and S. P. Pasca (2024). “Host circuit engagement of human cortical organoids transplanted in rodents.” Nat Protoc 19(12): 3542–67. [https://www.ncbi.nlm.nih.gov/pubmed/39075308]

Kettenmann, H., U. K. Hanisch, M. Noda and A. Verkhratsky (2011). “Physiology of microglia.” Physiol Rev 91(2): 461–553. [https://www.ncbi.nlm.nih.gov/pubmed/21527731]

Kitahara, T., H. Sakaguchi, A. Morizane, T. Kikuchi, S. Miyamoto and J. Takahashi (2020). “Axonal Extensions along Corticospinal Tracts from Transplanted Human Cerebral Organoids.” Stem Cell Reports 15(2): 467–81. [https://www.ncbi.nlm.nih.gov/pubmed/32679062]

Lancaster, M. A., M. Renner, C. A. Martin, D. Wenzel, L. S. Bicknell, M. E. Hurles, T. Homfray, J. M. Penninger, A. P. Jackson and J. A. Knoblich (2013). “Cerebral organoids model human brain development and microcephaly.” Nature 501(7467): 373–9. [https://www.ncbi.nlm.nih.gov/pubmed/23995685]

Li, Z., J. A. Klein, S. Rampam, R. Kurzion, N. B. Campbell, Y. Patel, T. F. Haydar and E. Zeldich (2022). “Asynchronous excitatory neuron development in an isogenic cortical spheroid model of Down syndrome.” Front Neurosci 16: 932384. [https://www.ncbi.nlm.nih.gov/pubmed/36161168]

Linaro, D., B. Vermaercke, R. Iwata, A. Ramaswamy, B. Libe-Philippot, L. Boubakar, B. A. Davis, K. Wierda, K. Davie, S. Poovathingal, P. A. Penttila, A. Bilheu, L. De Bruyne, D. Gall, K. K. Conzelmann, V. Bonin and P. Vanderhaeghen (2019). “Xenotransplanted Human Cortical Neurons Reveal Species-Specific Development and Functional Integration into Mouse Visual Circuits.” Neuron 104(5): 972–86 e6. [https://www.ncbi.nlm.nih.gov/pubmed/31761708]

Liu, X., Y. Lu, E. Iseri, C. Ren, H. Liu, T. Komiyama and D. Kuzum (2017). Transparent artifact-free graphene electrodes for compact closed-loop optogenetics systems. 2017 IEEE International Electron Devices Meeting (IEDM), IEEE.

Liu, X., C. Ren, Y. Lu, Y. Liu, J. H. Kim, S. Leutgeb, T. Komiyama and D. Kuzum (2021). “Multimodal neural recordings with Neuro-FITM uncover diverse patterns of cortical-hippocampal interactions.” Nat Neurosci 24(6): 886–96. [https://www.ncbi.nlm.nih.gov/pubmed/33875893]

Madhavan, M., Z. S. Nevin, H. E. Shick, E. Garrison, C. Clarkson-Paredes, M. Karl, B. L. L. Clayton, D. C. Factor, K. C. Allan, L. Barbar, T. Jain, P. Douvaras, V. Fossati, R. H. Miller and P. J. Tesar (2018). “Induction of myelinating oligodendrocytes in human cortical spheroids.” Nat Methods 15(9): 700–6. [https://www.ncbi.nlm.nih.gov/pubmed/30046099]

Mansour, A. A., J. T. Goncalves, C. W. Bloyd, H. Li, S. Fernandes, D. Quang, S. Johnston, S. L. Parylak, X. Jin and F. H. Gage (2018). “An in vivo model of functional and vascularized human brain organoids.” Nat Biotechnol 36(5): 432–41. [https://www.ncbi.nlm.nih.gov/pubmed/29658944]

Pachitariu, M., C. Stringer, M. Dipoppa, S. Schröder, L. F. Rossi, H. Dalgleish, M. Carandini and K. D. Harris (2017). “Suite2p: beyond 10,000 neurons with standard two-photon microscopy.” bioRxiv: 061507. [https://www.biorxiv.org/content/biorxiv/early/2017/07/20/061507.full.pdf]

Parr, C. J. C., S. Yamanaka and H. Saito (2017). “An update on stem cell biology and engineering for brain development.” Mol Psychiatry 22(6): 808–19. [https://www.ncbi.nlm.nih.gov/pubmed/28373686]

Pasca, A. M., S. A. Sloan, L. E. Clarke, Y. Tian, C. D. Makinson, N. Huber, C. H. Kim, J. Y. Park, N. A. O’Rourke, K. D. Nguyen, S. J. Smith, J. R. Huguenard, D. H. Geschwind, B. A. Barres and S. P. Pasca (2015). “Functional cortical neurons and astrocytes from human pluripotent stem cells in 3D culture.” Nat Methods 12(7): 671–8. [https://www.ncbi.nlm.nih.gov/pubmed/26005811]

Pires, J., R. Nelissen, H. D. Mansvelder and R. M. Meredith (2021). “Spontaneous synchronous network activity in the neonatal development of mPFC in mice.” Dev Neurobiol 81(2): 207–25. [https://www.ncbi.nlm.nih.gov/pubmed/33453138]

Pnevmatikakis, E. A. and A. Giovannucci (2017). “NoRMCorre: An online algorithm for piecewise rigid motion correction of calcium imaging data.” J Neurosci Methods 291: 83–94. [https://www.ncbi.nlm.nih.gov/pubmed/28782629]

Qian, X., H. Song and G. L. Ming (2019). “Brain organoids: advances, applications and challenges.” Development 146(8): dev166074. [https://www.ncbi.nlm.nih.gov/pubmed/30992274]

Ramezani, M., J. H. Kim, X. Liu, C. Ren, A. Alothman, C. De-Eknamkul, M. N. Wilson, E. Cubukcu, V. Gilja, T. Komiyama and D. Kuzum (2024). “High-density transparent graphene arrays for predicting cellular calcium activity at depth from surface potential recordings.” Nat Nanotechnol 19(4): 504–13. [https://www.ncbi.nlm.nih.gov/pubmed/38212523]

Revah, O., F. Gore, K. W. Kelley, J. Andersen, N. Sakai, X. Chen, M. Y. Li, F. Birey, X. Yang, N. L. Saw, S. W. Baker, N. D. Amin, S. Kulkarni, R. Mudipalli, B. Cui, S. Nishino, G. A. Grant, J. K. Knowles, M. Shamloo, J. R. Huguenard, K. Deisseroth and S. P. Pasca (2022). “Maturation and circuit integration of transplanted human cortical organoids.” Nature 610(7931): 319–26. [https://www.ncbi.nlm.nih.gov/pubmed/36224417]

Samarasinghe, R. A., O. A. Miranda, J. E. Buth, S. Mitchell, I. Ferando, M. Watanabe, T. F. Allison, A. Kurdian, N. N. Fotion, M. J. Gandal, P. Golshani, K. Plath, W. E. Lowry, J. M. Parent, I. Mody and B. G. Novitch (2021). “Identification of neural oscillations and epileptiform changes in human brain organoids.” Nat Neurosci 24(10): 1488–500. [https://www.ncbi.nlm.nih.gov/pubmed/34426698]

Schafer, S. T., A. A. Mansour, J. C. M. Schlachetzki, M. Pena, S. Ghassemzadeh, L. Mitchell, A. Mar, D. Quang, S. Stumpf, I. S. Ortiz, A. J. Lana, C. Baek, R. Zaghal, C. K. Glass, A. Nimmerjahn and F. H. Gage (2023). “An in vivo neuroimmune organoid model to study human microglia phenotypes.” Cell 186(10): 2111–26 e20. [https://www.ncbi.nlm.nih.gov/pubmed/37172564]

Shi, Y., L. Sun, M. Wang, J. Liu, S. Zhong, R. Li, P. Li, L. Guo, A. Fang, R. Chen, W. P. Ge, Q. Wu and X. Wang (2020). “Vascularized human cortical organoids (vOrganoids) model cortical development in vivo.” PLoS Biol 18(5): e3000705. [https://www.ncbi.nlm.nih.gov/pubmed/32401820]

Spitzer, N. C. (2006). “Electrical activity in early neuronal development.” Nature 444(7120): 707–12. [https://www.ncbi.nlm.nih.gov/pubmed/17151658]

Tang, J., X. Cheng, K. Kilic, A. Devor, J. Lee and D. A. Boas (2021). “Imaging localized fast optical signals of neural activation with optical coherence tomography in awake mice.” Opt Lett 46(7): 1744–7. [https://www.ncbi.nlm.nih.gov/pubmed/33793533]

Tang, R. P., S. Kelley, G. Kureli, E. A. Long, P. R. Bressan, S. Shah, S. E. Erdener, J. Jiang, J. T. Giblin, S. Kura, M. G. Simkulet, B. C. Rauscher, C. B. Schaffer, N. Nishimura, M. Thunemann, A. Devor and D. A. Boas (2025). “Capillaries susceptible to frequent stall dynamics revealed by comparing OCT and bessel-2PM measurements.” J Cereb Blood Flow Metab 45(10): 1905–17. [https://www.ncbi.nlm.nih.gov/pubmed/40562716]

Thunemann, M., Y. Lu, X. Liu, K. Kilic, M. Desjardins, M. Vandenberghe, S. Sadegh, P. A. Saisan, Q. Cheng, K. L. Weldy, H. Lyu, S. Djurovic, O. A. Andreassen, A. M. Dale, A. Devor and D. Kuzum (2018). “Deep 2-photon imaging and artifact-free optogenetics through transparent graphene microelectrode arrays.” Nat Commun 9(1): 2035. [https://www.ncbi.nlm.nih.gov/pubmed/29789548]

Trujillo, C. A., R. Gao, P. D. Negraes, J. Gu, J. Buchanan, S. Preissl, A. Wang, W. Wu, G. G. Haddad, I. A. Chaim, A. Domissy, M. Vandenberghe, A. Devor, G. W. Yeo, B. Voytek and A. R. Muotri (2019). “Complex Oscillatory Waves Emerging from Cortical Organoids Model Early Human Brain Network Development.” Cell Stem Cell 25(4): 558–69 e7. [https://www.ncbi.nlm.nih.gov/pubmed/31474560]

Voleti, V., K. B. Patel, W. Li, C. Perez Campos, S. Bharadwaj, H. Yu, C. Ford, M. J. Casper, R. W. Yan, W. Liang, C. Wen, K. D. Kimura, K. L. Targoff and E. M. C. Hillman (2019). “Real-time volumetric microscopy of in vivo dynamics and large-scale samples with SCAPE 2.0.” Nat Methods 16(10): 1054–62. [https://www.ncbi.nlm.nih.gov/pubmed/31562489]

Wang, D., X. Zhang, X. Y. Tang, Y. Gan, H. Yu, S. Wu, Y. Hong, M. Tao, C. Chu, X. Qi, H. Hu, Y. Zhu, W. Zhu, X. Han, M. Xu, Y. Dong, Q. Cheng, X. Guo and Y. Liu (2025). “Generation of human nucleus basalis organoids with functional nbM-cortical cholinergic projections in transplanted assembloids.” Cell Stem Cell 32(12): 1833–48 e7. [https://www.ncbi.nlm.nih.gov/pubmed/41187744]

Wang, J., G. Qiao, H. Tan, M. J. Li, K. Wang, J. Xu, M. Janowski, T.-M. Fu, P. Walczak and Y. Liang (2025). “Intravital Multimodal Imaging of Human Cortical Organoids for Chronic Stroke Treatment in Mice.” bioRxiv: 2025.07.05.663297. [https://www.biorxiv.org/content/biorxiv/early/2025/07/06/2025.07.05.663297.full.pdf]

Wang, R. K., Q. Zhang, Y. Li and S. Song (2017). “Optical coherence tomography angiography-based capillary velocimetry.” J Biomed Opt 22(6): 66008. [https://www.ncbi.nlm.nih.gov/pubmed/28617921]

Wang, Y., Y. Zheng, X. Xu, E. Dubuisson, Q. Bao, J. Lu and K. P. Loh (2011). “Electrochemical delamination of CVD-grown graphene film: toward the recyclable use of copper catalyst.” ACS Nano 5(12): 9927–33. [https://www.ncbi.nlm.nih.gov/pubmed/22034835]

Warm, D., D. Bassetti, L. Gellert, J. W. Yang, H. J. Luhmann and A. Sinning (2025). “Spontaneous mesoscale calcium dynamics reflect the development of the modular functional architecture of the mouse cerebral cortex.” Neuroimage 309: 121088. [https://www.ncbi.nlm.nih.gov/pubmed/39954874]

Watanabe, M., J. E. Buth, N. Vishlaghi, L. de la Torre-Ubieta, J. Taxidis, B. S. Khakh, G. Coppola, C. A. Pearson, K. Yamauchi, D. Gong, X. Dai, R. Damoiseaux, R. Aliyari, S. Liebscher, K. Schenke-Layland, C. Caneda, E. J. Huang, Y. Zhang, G. Cheng, D. H. Geschwind, P. Golshani, R. Sun and B. G. Novitch (2017). “Self-Organized Cerebral Organoids with Human-Specific Features Predict Effective Drugs to Combat Zika Virus Infection.” Cell Rep 21(2): 517–32. [https://www.ncbi.nlm.nih.gov/pubmed/29020636]

Wilson, M. N., M. Thunemann, X. Liu, Y. Lu, F. Puppo, J. W. Adams, J. H. Kim, M. Ramezani, D. P. Pizzo, S. Djurovic, O. A. Andreassen, A. A. Mansour, F. H. Gage, A. R. Muotri, A. Devor and D. Kuzum (2022). “Multimodal monitoring of human cortical organoids implanted in mice reveal functional connection with visual cortex.” Nat Commun 13(1): 7945. [https://www.ncbi.nlm.nih.gov/pubmed/36572698]

Wonders, C. P. and S. A. Anderson (2006). “The origin and specification of cortical interneurons.” Nat Rev Neurosci 7(9): 687–96. [https://www.ncbi.nlm.nih.gov/pubmed/16883309]

Wu, M. W., N. Kourdougli and C. Portera-Cailliau (2024). “Network state transitions during cortical development.” Nat Rev Neurosci 25(8): 535–52. [https://www.ncbi.nlm.nih.gov/pubmed/38783147]

Yao, P., R. Liu, T. Broggini, M. Thunemann and D. Kleinfeld (2023). “Construction and use of an adaptive optics two-photon microscope with direct wavefront sensing.” Nat Protoc 18(12): 3732–66. [https://www.ncbi.nlm.nih.gov/pubmed/37914781]

Zhang, W., J. Jiang, Z. Xu, H. Yan, B. Tang, C. Liu, C. Chen and Q. Meng (2023). “Microglia-containing human brain organoids for the study of brain development and pathology.” Mol Psychiatry 28(1): 96–107. [https://www.ncbi.nlm.nih.gov/pubmed/36474001]

Zheng, C. X., S. M. Wang, Y. H. Bai, T. T. Luo, J. Q. Wang, C. Q. Dai, B. L. Guo, S. C. Luo, D. H. Wang, Y. L. Yang and Y. Y. Wang (2018). “Lentiviral Vectors and Adeno-Associated Virus Vectors: Useful Tools for Gene Transfer in Pain Research.” Anat Rec (Hoboken) 301(5): 825–36. [https://www.ncbi.nlm.nih.gov/pubmed/29149775]

Zudaire, E., L. Gambardella, C. Kurcz and S. Vermeren (2011). “A computational tool for quantitative analysis of vascular networks.” PLoS One 6(11): e27385. [https://www.ncbi.nlm.nih.gov/pubmed/22110636]

