## Supplemental Material for "A neurorecording toolkit for longitudinal assessments of transplanted human cortical organoids *in vivo*"

### Supplementary Tables

**Supplementary Table 1. Study parameters per experimental animal.** From 36 animals, 11 animals died peri-operatively (1) or due to complications within the first 0-8 post-operative days (10); 3 animals were found dead 51-233 days after surgery; 22 animals were euthanized 50-294 days after xenograft implantation for ex vivo analyses.

| ID | Batch | Protocol | Cell Line | Transduction | Array | Follow-up [days] | Cause of death | Data |
|---|---|---|---|---|---|---|---|---|
| 001 | a | A | WT83(Tang *et al.* 2016) | AAV7m8 hSyn1-GCaMP6s-p2A-NLS-tdTomato | - | 2 | Died during recovery | Figures 1 (IHC), 7, S1, S8 |
| 002 | | | | | 16-ch | 5 | Died during recovery | |
| 003 | | | | | 16-ch | 131 | Found dead | |
| 004 | | | | | 16-ch | 1 | Died during recovery | |
| 005 | | | | | 16-ch | 293 | Tissue extraction | |
| 006 | | | | | 16-ch | 293 | Tissue extraction | |
| 007 | | | | | 16-ch | 292 | Tissue extraction | |
| 008 | | | | | 16-ch | 292 | Tissue extraction | |
| 009 | | | | | 16-ch | 8 | Died during recovery | |
| 010 | | | | | - | 0 | Peri-operatively | |
| 011 | | | | | 16-ch | 290 | Tissue extraction | |
| 012 | | | | | 16-ch | 233 | Found dead | |
| 013 | b | B | WC-24-02-DS-B | pGP-AAV1-synjGCaMP8s-WPRE<br><br>pAAV8-CAG-tdTomato (codon diversified) | - | 167 | Tissue extraction | Figure 1 (brightfield), Video S1 |
| 014 | | | | | - | 163 | Tissue extraction | |
| 015 | c | | | | - | 97 | Tissue extraction | Figures 3 (AAV data – 2P and IHC), S5 (AAV) |
| 016 | | | | | - | 1 | Died during recovery | |
| 017 | | | | | - | 94 | Tissue extraction | |
| 018 | | | | | - | 93 | Tissue extraction | |
| 019 | | | | | - | 0 | Died during recovery | |
| 020 | | | | | - | 4 | Died during recovery | |
| 021 | | | | | - | 50 | Tissue extraction | |
| 022 | | | | | - | 3 | Died during recovery | |
| 023 | | | | | - | 8 | Died during recovery | |
| 024 | | | | | - | 51 | Found dead | |
| 025 | d | | | pLV[Exp]-Puro-SYN1.jGCaMP8s<br><br>pLV[Exp]-Bsd-EF1A>mScarlet3 | - | 103 | Tissue extraction | Figures 3 (LV data – 2P and IHC), 4, 5, S5 (LV), S6, S7, Videos S2-4 |
| 026 | | | | | - | 102 | Tissue extraction | |
| 027 | | | | | - | 102 | Tissue extraction | |
| 028 | | | | | - | 1 | Died during recovery | |
| 029 | | | | | - | 107 | Tissue extraction | |
| 030 | | | | | - | 101 | Tissue extraction | |
| 031 | e | | | | - | 154 | Tissue extraction | Figures 2, S3, S4 |
| 032 | | | WC-24-02-DS-M | | - | 161 | Tissue extraction | |
| 033 | | | | | - | 153 | Tissue extraction | |
| 034 | | | | | - | 157 | Tissue extraction | |
| 035 | f | | MGH2046 (Seo *et al.* 2017) | | - | 294 | Tissue extraction | Figure 6 |
| 036 | | | | | - | 140 | Tissue extraction | |

### Supplementary Figures

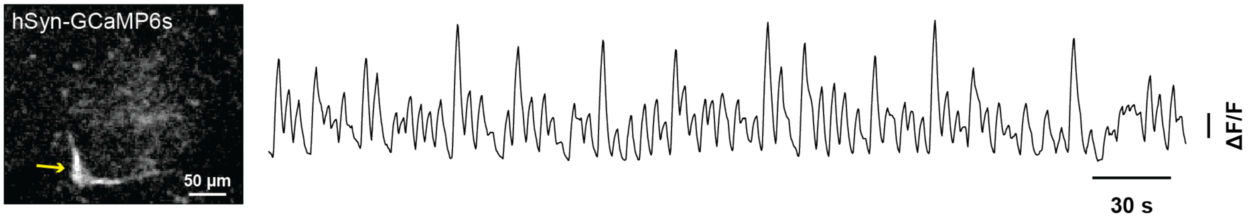


**Supplementary Figure 1. *In vivo* two-photon calcium imaging of human cortical organoid (hCO) xenograft neuron eight months after xenotransplantation.** The yellow arrow points towards a functionally active, GCaMP6s-expressing, hCO-derived neuron; the corresponding activity trace (as baseline-normalized ΔF/F) is shown on the right. The hCO was generated by Protocol A (Trujillo *et al.* 2019) and transduced in culture using AAV7m8 hSyn1-GCaMP6s-p2A-NLS-tdTomato.

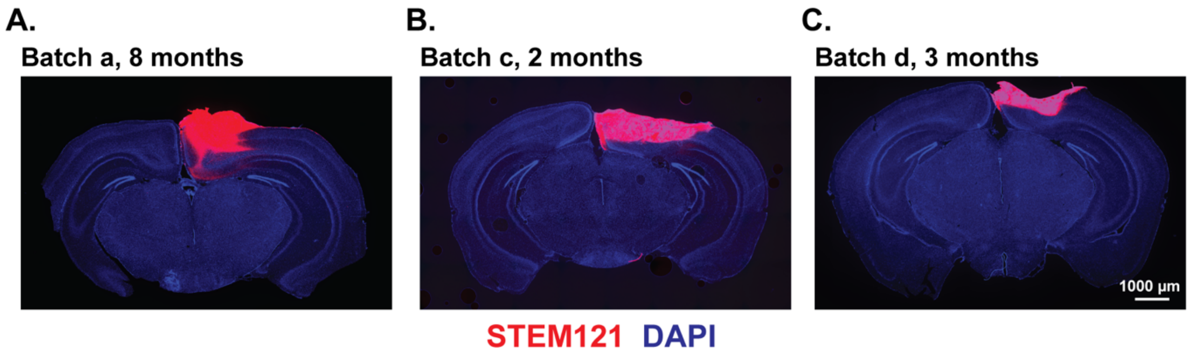


**Supplementary Figure 2. Brain tissue sections with xenotransplanted hCOs show successful engraftment of hCOs generated with two different protocols.** Immunostaining for human cytoplasm (STEM121) shows hCO xenograft borders within coronal sections of the host brain. Organoids were derived from batches as denoted in **Supplementary Table 1**.

A. Batch a (Trujillo *et al.* 2019), 8 months after xenotransplantation.
B. Batch c (Madhavan *et al.* 2018), 2 months after xenotransplantation.
C. Batch d (Madhavan *et al.* 2018), 3 months after xenotransplantation.

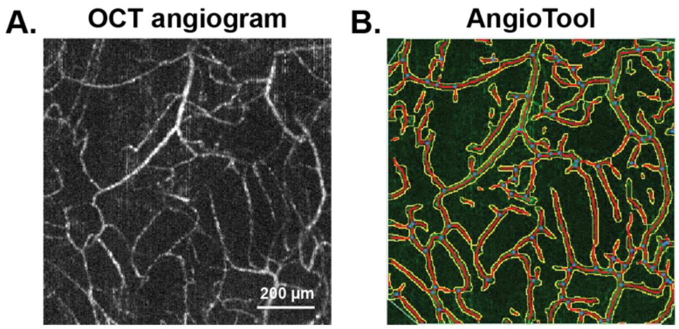


**Supplementary Figure 3. Quantification of capillary density using automated vessel detection through AngioTool (Zudaire *et al.* 2011).**

A. 500-µm maximum intensity projection across the Z axis ($MIP_Z$) angiogram taken with optical coherence tomography (OCT) showing capillaries within the xenograft tissue two months after xenotransplantation.
B. Overlay from Angiotool to quantify capillary density. Yellow lines outline capillaries, red lines show capillary skeleton, and blue dots show branching points.

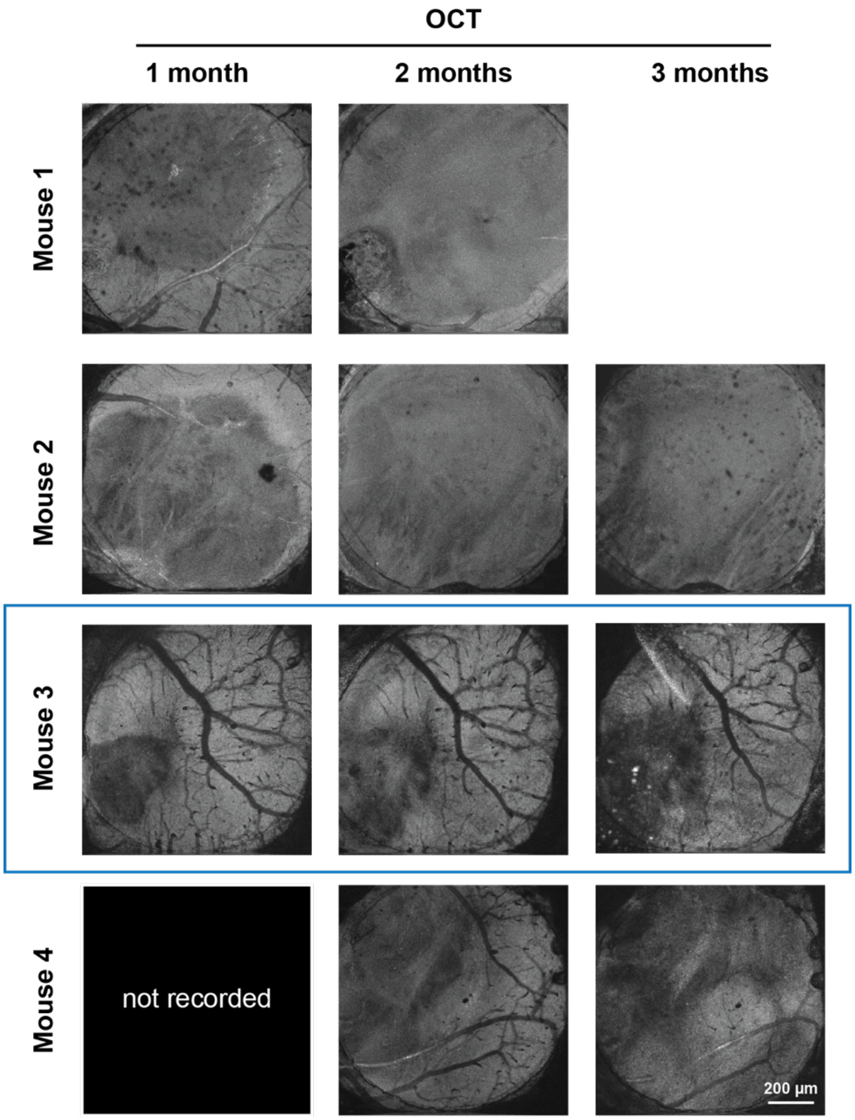


**Supplementary Figure 4. Optical coherence tomography (OCT) angiograms taken at 1, 2, and 3 months after xenotransplantation.** OCT 100-µm maximum intensity projection along the Z axis ($MIP_Z$) for four animals. All xenografts recorded at one month were visible within the optical window. At 2-3 months, most xenografts had grown to exceed the bounds of the optical window. The xenograft in Mouse 3 (blue box) is shown and quantified in **Figure 2** as it was the only xenograft that remained entirely in the field of view at all imaging time points.

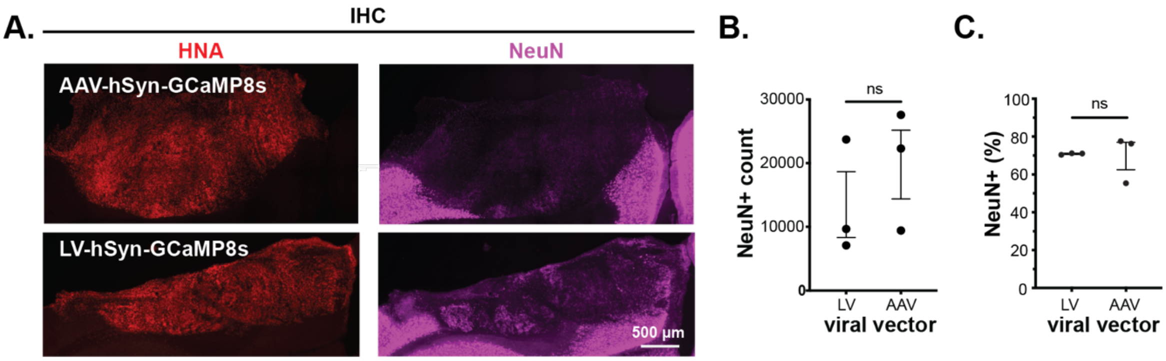


**Supplementary Figure 5. Immunostaining for NeuN shows no significant difference in neuron density between AAV- and LV-labelled xenografts in vivo.**

A. Immunostaining for human nuclei (HNA) and neurons (NeuN) for hCO xenografts labelled using either AAV-hSyn-GCaMP8s (top) or LV-hSyn-GCaMP8s (bottom) vectors. Note that in our hands, NeuN staining tends to be weaker for human than for mouse neurons (Wilson *et al.* 2022).

B. Quantification of cell counts for NeuN-positive (NeuN+) cells within the graft showing that there are no statistical differences in neuronal density between AAV- and LV-labelled hCO xenografts at 3 months post-xenotransplantation. ns = not significant; p=0.4478 (two-tailed t-test). Plot shows mean ± s.e.m. for N=3 mice per condition.

C. Quantification of percent NeuN+ cells of total HNA+ cells showing that there are no statistical differences in proportion of neurons to overall cell counts within AAV- and LV- labelled hCO xenografts at three months post-xenotransplantation. NS = not significant; p=0.8889 (two-tailed t-test). Bar chart shows mean ± s.e.m. for N=3 mice per condition.

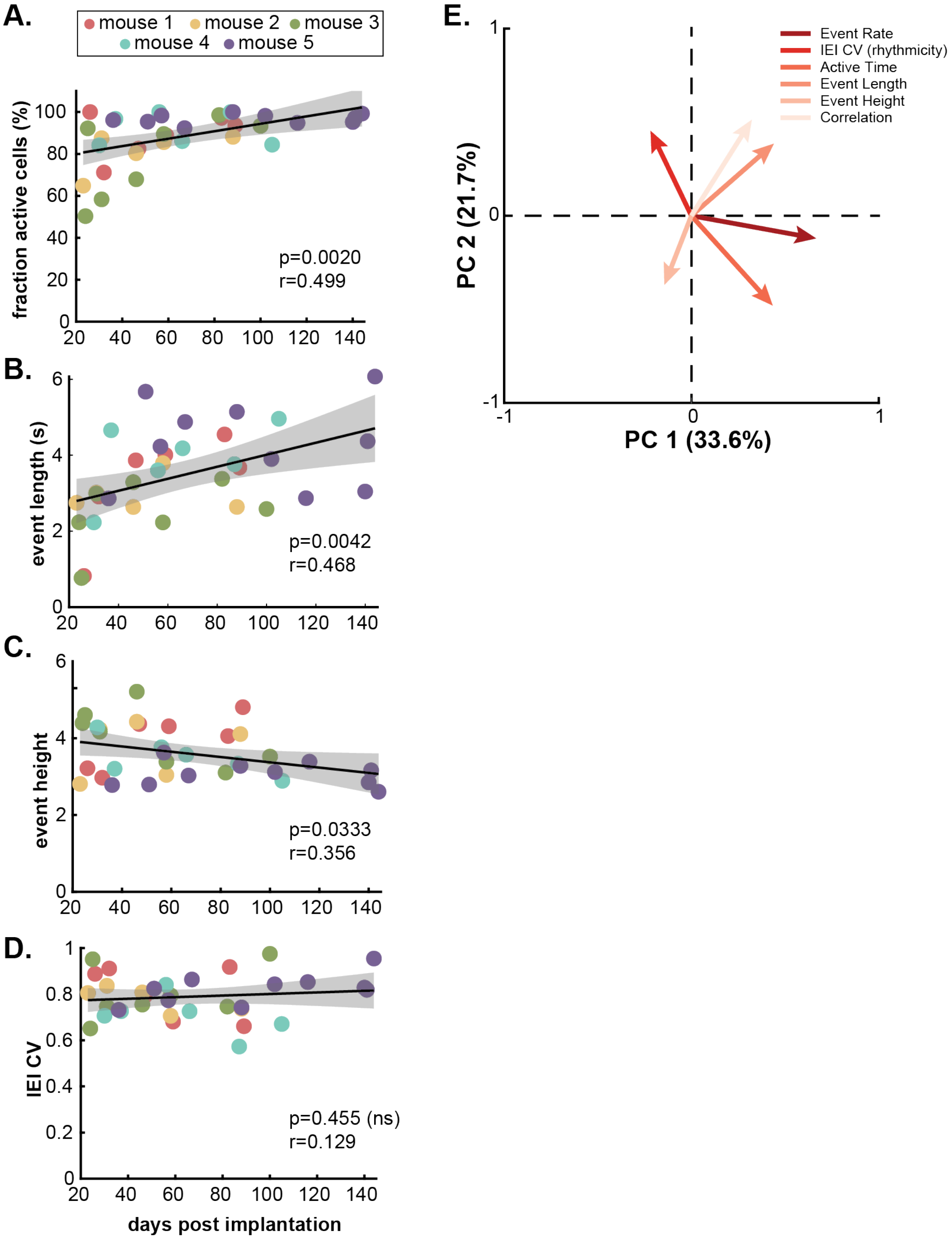


**Supplementary Figure 6. Longitudinal trends in neuronal activity of xenotransplanted hCO neurons measured via calcium imaging.**

A. Longitudinal changes in the fraction of active cells over 3 months for each animal and imaging session. Active cells were defined as having at least one calcium event over a 3-5-min recording period (N=5 animals). Scatterplot shows linear mixed effects model (LMM; p=0.0020, r=0.499).
B. Longitudinal changes in calcium event length (seconds) over 3 months for each animal and imaging session (N=5 animals). Scatterplot shows linear mixed effects model (LMM; p=0.0042, r=0.468).
C. Longitudinal changes in calcium event height (baseline-normalized ΔF/F) over 3 months for each animal and imaging session (N=5 animals). Scatterplot shows linear mixed effects model (LMM; p=0.0020, r=0.499).
D. Longitudinal changes in coefficient of variations (CV) for the inter-event-interval (IEI) over 3 months for each animal and imaging session (N=5 animals). Scatterplot shows linear mixed effects model (LMM; p=0.455, ns=not significant, r=0.129).
E. Feature weights for the PCA shown in **Figure 4H-I**.

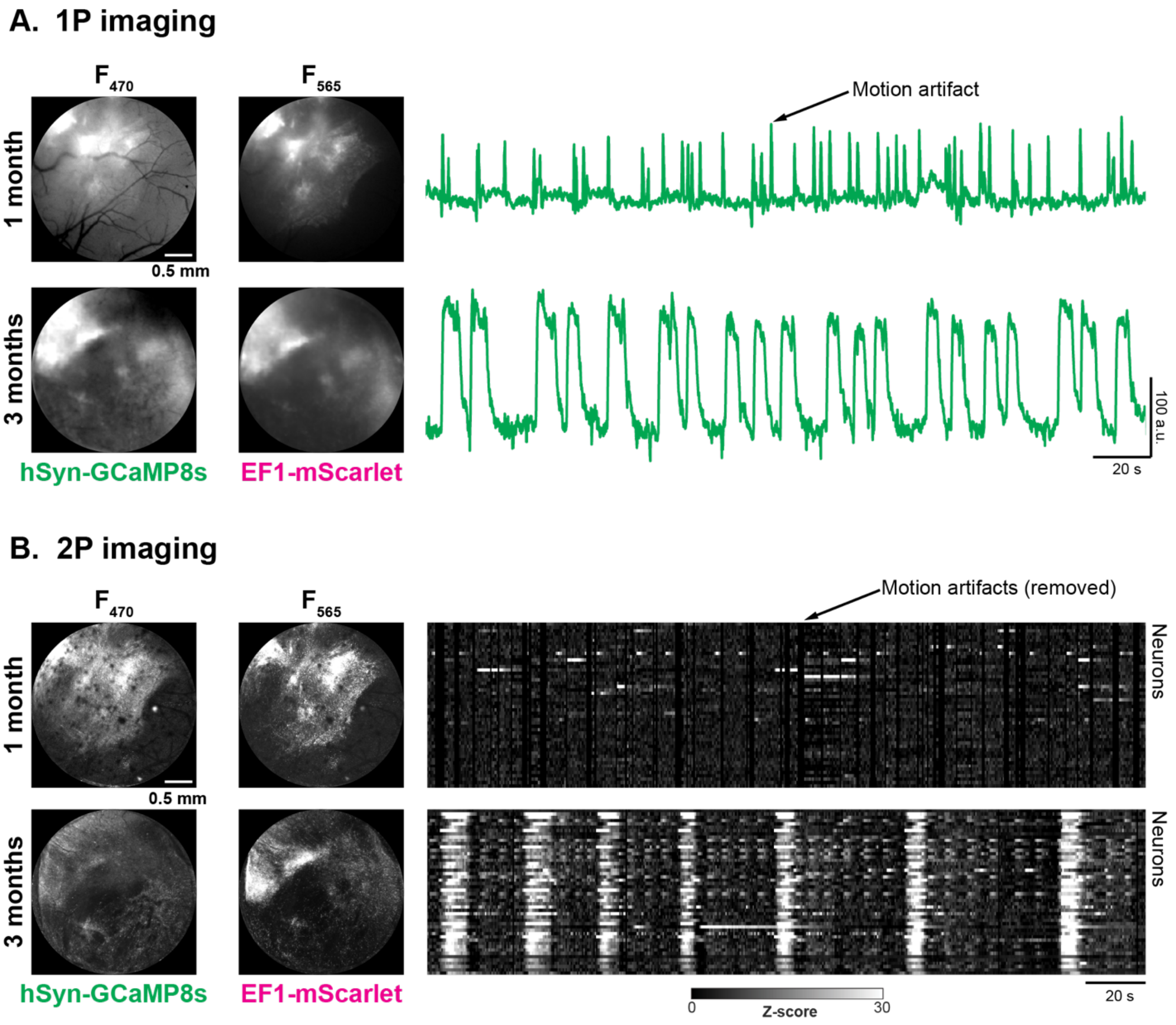


**Supplementary Figure 7. One-photon widefield calcium imaging shows transition towards synchronized hCO xenograft activity.**

A. One-photon (1P) imaging in the same animal one and three months after xenotransplantation; (left) overview images of the exposure acquired as fluorescence at 470-nm (GCaMP8s) and 565-nm (mScarlet) excitation; (right) time courses of ($F_{565}$-normalized) $F_{470}$ signal changes recorded within a region of interest covering the xenograft. Overview images show expansion of the xenograft over the 2-month period. Note motion artifacts in 1-month recording appearing as transient spikes within the normalized $F_{470}$*-signal that are not being removed by image registration.

B. Two-photon (2P) imaging of the same animal as shown in panel A one and three months after xenotransplantation; (left) low-magnification maximum intensity projections of Z-stacks of the entire exposure; (right) heat map showing changes in GCaMP8s fluorescence (shown as baseline-normalized Z-score) of individual neurons from representative time-series recordings. Note the increased incidence of motion artifacts one month after transplantation; here, periods of excessive motion are set to 'NaN'.

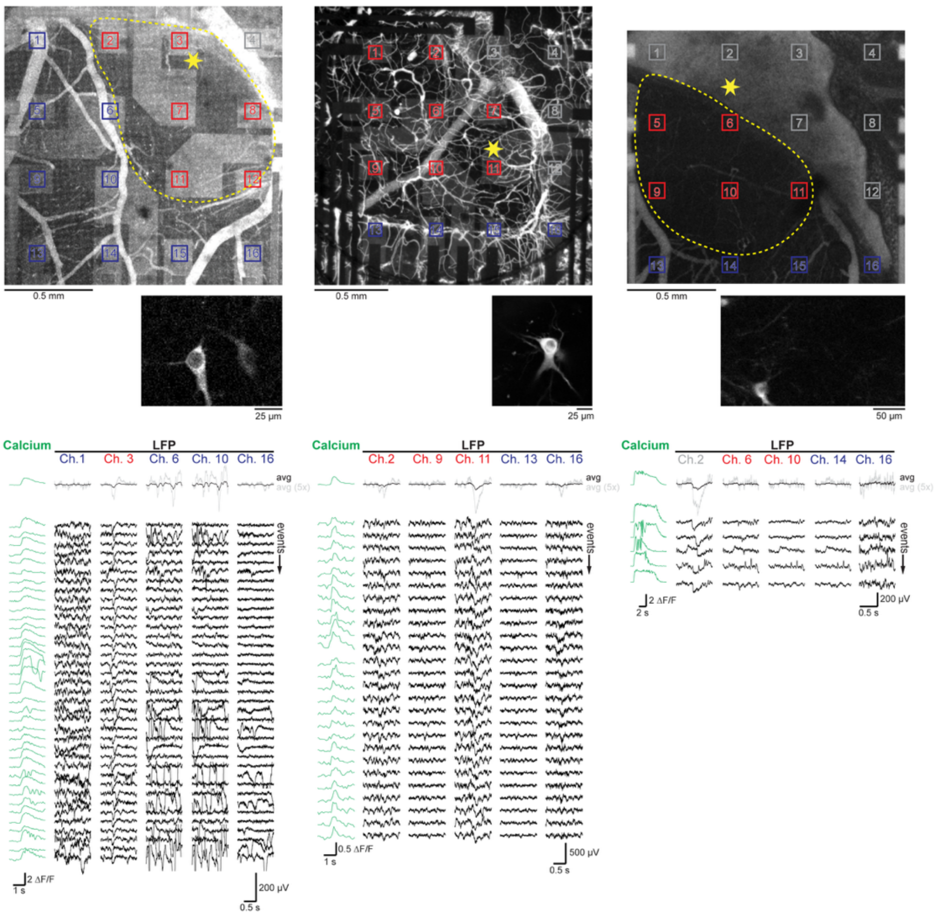


**Supplementary Figure 8. Electrocorticography recordings of local field potential (LFP) with graphene microelectrode arrays (gMEAs) correspond to calcium events in human neurons.**

(Top) Overview of the exposure acquired with two-photon (2P) imaging after labeling the blood plasma with Alexa680-Dextran for 3 animals. Xenotransplant boundaries are indicated by the yellow dotted line. Graphene electrodes are highlighted in blue, red, and grey boxes corresponding to their location above host cortex, xenograft, and bone, respectively. The yellow star indicates the location where 2P imaging was performed. (Middle) 2P image from the field-of-view of the respective calcium imaging timeseries. (Bottom) Calcium-event triggered average (avg) and individual calcium events and corresponding local field potential (LFP) signals from selected channels across the gMEA. LFP signals are temporally aligned with individual calcium events. (For display, channels with impedances <5 MΩ were chosen). The left was taken from the same animal and recording session as the data shown in **Figure 7** (a different GCaMP6s-expressing neuron was chosen); for the right, the 2P vascular map (top) was taken ~4 weeks after functional recordings (middle/bottom), during which significant bone regrowth (as seen in the top left part of the image on the top), now covering channel 2, occurred.

#### Supplementary Videos

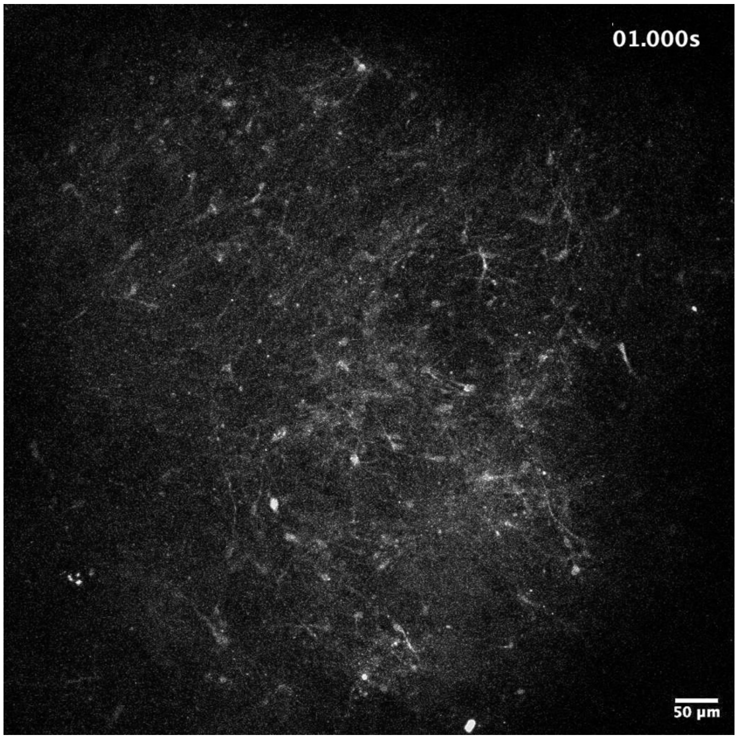


**Supplementary Video 1. *In vitro* calcium imaging of an AAV-labelled hCO.** We used AAV to express GCaMP8s in hCOs under the human synapsin (hSyn) promoter leading to sufficient labelling density to visualize neuronal activity at 120 days *in vitro* (DIV). In the example shown, the organoid is labelled at 105 days DIV using AAV1-hSyn-GCaMP8s. The video is taken at 120 DIV using a Zeiss LSM 710-Live Duo Confocal with 10x (NA=0.45) objective and incubator attachment (5% $CO_2$ and 37 °C) at 1 Hz for 180 s; video presented at 10 frames per second.

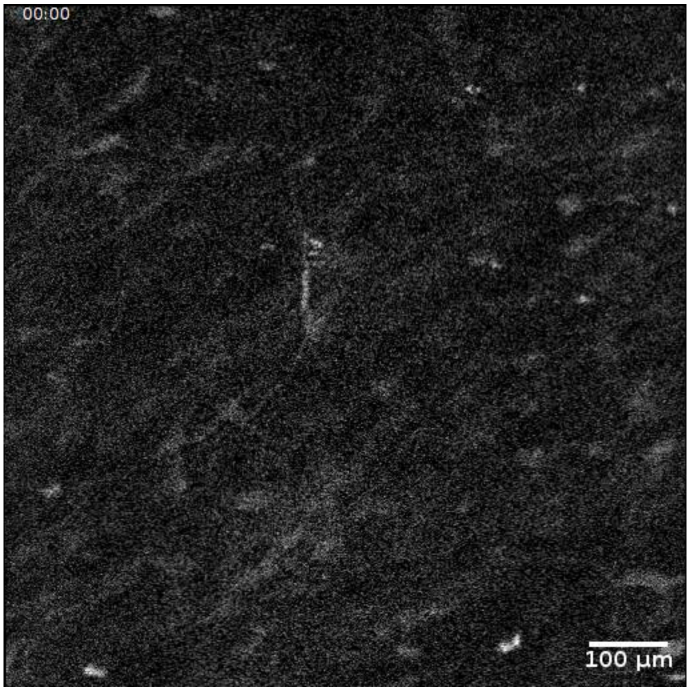


**Supplementary Video 2. *In vivo* two-photon calcium imaging of hCO neurons at one month after xenotransplantation.** Here, we used LV to express GCaMP8s in hCOs under the human synapsin (hSyn) promoter and performed xenotransplantation after 40 days *in vitro* (DIV). In the example shown, the video is taken one month after xenotransplantation.

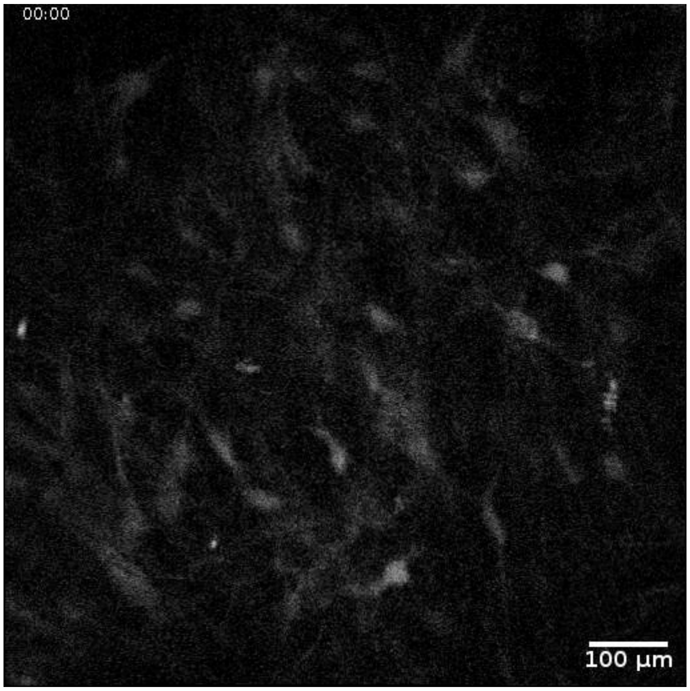


**Supplementary Video 3. *In vivo* two-photon calcium imaging of hCO neurons at three months after xenotransplantation.** Here, we used LV to express GCaMP8s in hCOs under the human synapsin (hSyn) promoter and performed xenotransplantation after 40 days *in vitro* (DIV). In the example shown, the video is taken three months after xenotransplantation.

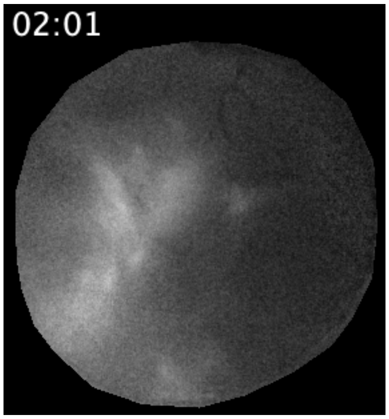


**Supplementary Video 4. One-photon widefield calcium imaging at three months after xenotransplantation.** The video corresponds to run 05 in **Figure 5** and shows the $F_{470}$* time course, i.e., the change in GCaMP8s fluorescence at 470-nm excitation corrected by (mScarlet) fluorescence at 565-nm excitation. The frame rate was reduced from 15 Hz to 3 Hz by averaging five consecutive frames; the final video shows the time course at 5x of the original speed. The field-of-view corresponds to 2.9 mm x 3.1mm; areas surrounding the cranial window are being masked and set to 'NaN'.

### Supplementary References

Madhavan, M., Z. S. Nevin, H. E. Shick, E. Garrison, C. Clarkson-Paredes, M. Karl, B. L. L. Clayton, D. C. Factor, K. C. Allan, L. Barbar, T. Jain, P. Douvaras, V. Fossati, R. H. Miller and P. J. Tesar (2018). "Induction of myelinating oligodendrocytes in human cortical spheroids." Nature Methods **15**(9): 700–6.

Seo, J., O. Kritskiy, L. A. Watson, S. J. Barker, D. Dey, W. K. Raja, Y.-T. Lin, T. Ko, S. Cho, J. Penney, M. C. Silva, S. D. Sheridan, D. Lucente, J. F. Gusella, B. C. Dickerson, S. J. Haggarty and L.-H. Tsai (2017). "Inhibition of p25/Cdk5 Attenuates Tauopathy in Mouse and iPSC Models of Frontotemporal Dementia." The Journal of Neuroscience **37**(41): 9917–24.

Tang, X., J. Kim, L. Zhou, E. Wengert, L. Zhang, Z. Wu, C. Carromeu, A. R. Muotri, M. C. N. Marchetto, F. H. Gage and G. Chen (2016). "KCC2 rescues functional deficits in human neurons derived from patients with Rett syndrome." Proceedings of the National Academy of Sciences **113**(3): 751–6.

Trujillo, C. A., R. Gao, P. D. Negraes, J. Gu, J. Buchanan, S. Preissl, A. Wang, W. Wu, G. G. Haddad, I. A. Chaim, A. Domissy, M. Vandenberghe, A. Devor, G. W. Yeo, B. Voytek and A. R. Muotri (2019). "Complex Oscillatory Waves Emerging from Cortical Organoids Model Early Human Brain Network Development." Cell Stem Cell **25**(4): 558–69.e7.

Wilson, M. N., M. Thunemann, X. Liu, Y. Lu, F. Puppo, J. W. Adams, J. H. Kim, M. Ramezani, D. P. Pizzo, S. Djurovic, O. A. Andreassen, A. A. Mansour, F. H. Gage, A. R. Muotri, A. Devor and D. Kuzum (2022). "Multimodal monitoring of human cortical organoids implanted in mice reveal functional connection with visual cortex." Nat Commun **13**(1): 7945.

Zudaire, E., L. Gambardella, C. Kurcz and S. Vermeren (2011). "A computational tool for quantitative analysis of vascular networks." PLoS One **6**(11): e27385.
